# The recoverable resolution of cellular perturbation-response prediction

**DOI:** 10.64898/2026.09.30.755756

**Authors:** Yongqi Huang, Hanzhi Wang, Chenyu Li, Parker Wilson

## Abstract

Virtual cell models aim to predict how cells respond to perturbations, yet accurate prediction of post-perturbation gene expression can conceal failures to recover context-specific responses. Here we identify context geometry compression (CGC), in which models preserve cellular identity and shared perturbation effects while collapsing differences in how the same perturbation acts across contexts. Across biological systems, reproducible context-specific responses remained difficult to recover; in a large drug-response matrix, this persisted even with nearly complete perturbation coverage. The same drug-response predictions achieved higher normalized recovery in predefined pathway summaries than in gene-level response profiles, while coarse response programs were more predictable in human lymphoblastoid cell lines. Compact baseline transcriptomic representations retained most full-RNA predictive performance for patient-specific ex vivo drug responses. Together, these findings establish recoverable biological resolution, rather than measurement dimensionality, as a criterion for evaluating virtual cell predictions and matching model outputs to the biological detail supported by evidence.

## Introduction

The central goal of virtual cell models is to predict how cells respond to perturbations^1,2^. Yet models are often asked to predict perturbation responses across thousands of genes using a limited snapshot of the baseline cell state, typically a gene-expression profile^3–6^. A transcriptomic snapshot may accurately describe a cell’s current state without fully capturing the regulatory information that shapes its response to perturbation.

Most perturbation-prediction benchmarks nevertheless evaluate the post-perturbation state directly^3–6^. For perturbation *p* in cellular context *c*, we write the perturbed state as *Y*_*p,c*_ =*X*_*c*_ + *Δ*_*p,c*_. The baseline state *X*_*c*_ can dominate similarity between prediction and observation even when the perturbation response *Δ*_*p,c*_ is poorly recovered. Recent studies support this concern, showing that apparent gains in perturbation prediction can diminish when models are evaluated against simple baselines or with metrics that isolate perturbation-specific effects^7,8^. This raises a central question: after accounting for baseline state and shared perturbation effects, how much of the context-specific response can a model recover?

In our complementary work on generalization to unseen perturbations, we observed intervention geometry compression (IGC), in which models preserved broad perturbational structure while collapsing the geometry that distinguishes unseen perturbations^9^. Here we examine the complementary problem of generalization across cellular contexts: can a model recover how the same perturbation acts differently across biological contexts?

We decompose each perturbation response as *Δ*_*p,c*_ = *Δ*_*p*_ + *Γ*_*p,c*_, where *Δ*_*p*_ is the response shared across contexts and *Γ*_*p,c*_ is the context-specific component. For a given context, the mapping from a perturbation to this component defines its context-specific response operator. We call the dissociation between recognizing a context and recovering this response rule the identity–operator gap.

Across primary human T cells^10^, a contemporary zero-shot cytokine model^6,11^, a large replicated drug-response matrix^12^ and human lymphoblastoid cell lines^13^, we identify context geometry compression (CGC), in which models preserve cellular identity and shared perturbation effects while compressing differences in responses across contexts. In a prospective completion test of a 50-context × 93-perturbation matrix^12^, context-specific recovery became detectable only after more than 80% of entries were observed and remained 0.111 even when 4,649 of 4,650 entries had been measured. Baseline transcriptomic state supported prediction of coarse response programs even when individual-specific gene-level responses remained nearly unrecovered. Residual responses pooled across context–perturbation pairs remained high-dimensional, yet projecting the same predictions into predefined pathways yielded higher normalized recovery. In patient-derived organoids^14^, drug-response prediction performance saturated within 16–32 transcriptomic components derived from full RNA, despite 19,421 measured genes.

Together, these analyses establish a resolution-aware framework for cellular perturbation prediction. The framework separates the detail measured by an assay, the detail that can be predicted from the available information and the detail requested of a model. We define recoverable resolution as the level of biological detail at which responses can be reproducibly predicted from information available at prediction time. This makes biological resolution an explicit criterion for evaluating virtual cells and the experiments they are intended to replace. It provides a basis for choosing which response features to predict and testing how much experimental support those predictions require.

## Results

### State-level accuracy conceals context geometry compression

We first asked whether high post-perturbation state accuracy necessarily implies recovery of the context-specific response. In a primary human CD4+ T-cell CRISPRi Perturb-seq dataset^10^, donor identity defined the biological context and Rest, 8-h stimulation and 48-h stimulation provided within-donor states. We framed the prediction problem as an intervention × context response surface (Fig. 1a), then used guide-disjoint response halves and replicate-disjoint comparisons to isolate reproducible stimulation-dependent operator structure before evaluating any predictive model (Fig. 1b,c).

**Figure 1.**
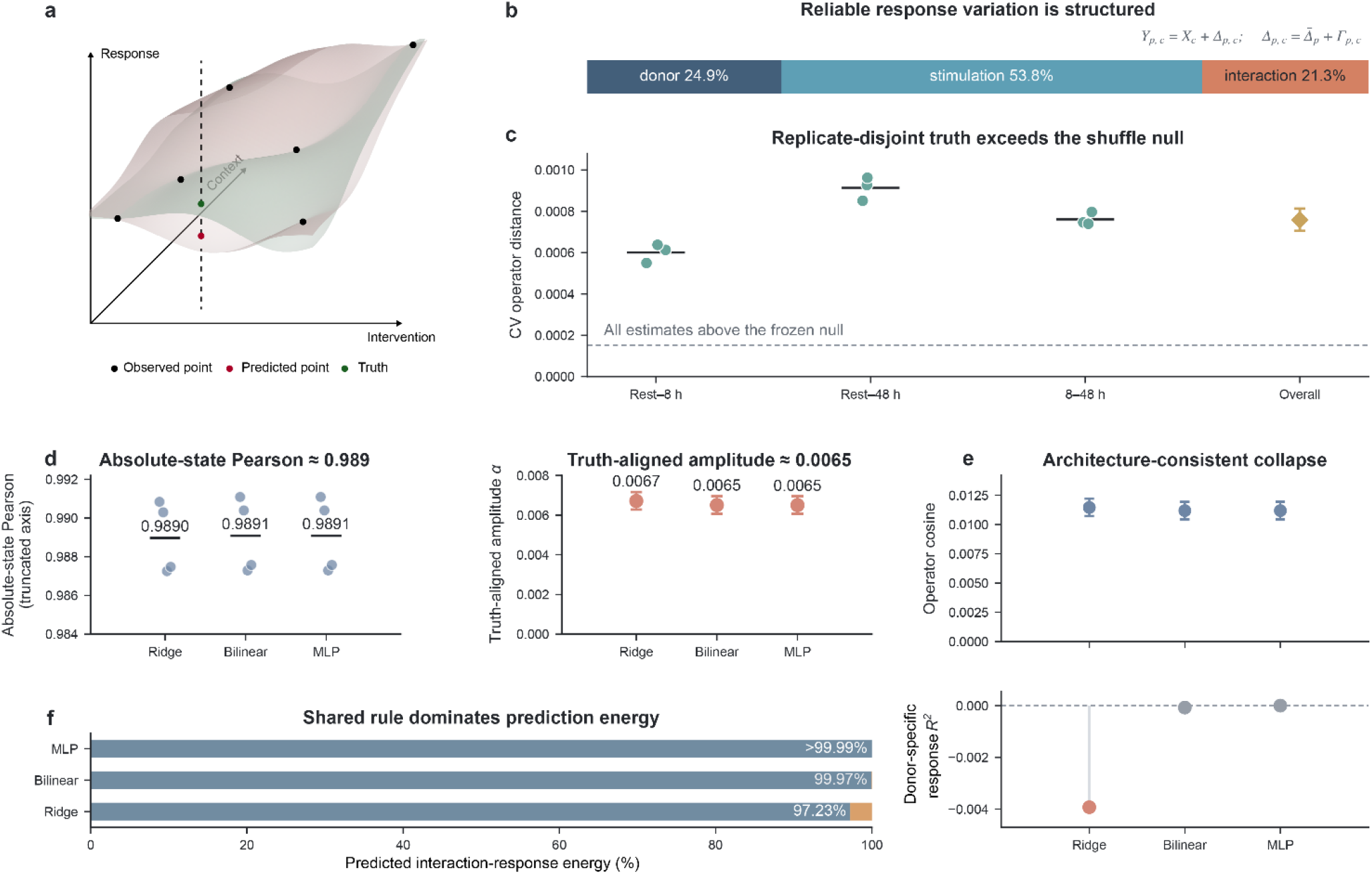
State-level accuracy conceals context geometry compression. a, Conceptual intervention × context × response geometry; the schematic contains no numerical result. b, Positive-truncated, normalized point-estimate shares of reliable cross-guide energy attributed to intervention × donor (24.86%), intervention × stimulation (53.84%) and intervention × donor × stimulation (21.30%). These shares are distinct from bootstrap energy intervals; the intervention × donor component had bootstrap probability positive 0.9646. c, Replicate-disjoint truth gate across three donor partitions and three stimulation contrasts (n = 9 partition × contrast estimates). Overall operator distance was 0.000758 (95% bootstrap interval, 0.000705–0.000813), compared with the 95th percentile of the prespecified frozen shuffle null (0.000152; empirical P = 1/5001). d, Conditional absolute-state Pearson correlation and truth-aligned response amplitude for Ridge, bilinear and MLP models under strict leave-one-donor-out evaluation (n = 4 donors per model). e, Truth-aligned operator cosine and donor-specific response R^2^ for the same predictions. f, Fraction of predicted perturbation-by-stimulation interaction energy assigned by orthogonal projection to the training-donor shared response: 97.23%, 99.97% and >99.99%, respectively.

Pseudobulk response construction, guide-disjoint truth halves and strict held-donor information sets are detailed in Supplementary Fig. 1a–d. The cross-validated truth statistic was 7.58 × 10^−4^ (95% bootstrap interval, 7.05 × 10^−4^ −8.13 × 10^−4^), more than fourfold above the 95th percentile of the prespecified shuffle null (1.52 × 10^−4^; empirical *P* = 1/5001). Thus, context-specific response variation was experimentally reproducible rather than an artifact of measurement noise.

The apparent difficulty emerged only after separating absolute state from response geometry. Under strict leave-one-donor-out evaluation, Ridge, bilinear and multilayer-perceptron predictors retained conditional post-perturbation state Pearson correlations of approximately 0.989, yet truth-aligned context-specific response amplitude was only 0.00671, 0.00650 and 0.00650, respectively (Fig. 1d). Operator cosine was similarly small (0.0115, 0.0112 and 0.0112), and donor-specific response *R*^2^ was non-positive for all three model families (Fig. 1e). These metrics quantify alignment with the reproducible held-donor operator, rather than the fraction of biological variation present.

The predicted responses instead collapsed toward a shared rule learned from the training donors. The fraction of predicted perturbation-by-stimulation interaction energy aligned with the training-donor shared response was 97.23% for Ridge, 99.97% for the bilinear model and >99.99% for the MLP (Fig. 1f). Thus, near-perfect state-level prediction could coexist with near-complete loss of context-specific counterfactual amplitude. We refer to this systematic contraction of context-dependent response geometry as CGC.

### Context identity does not guarantee stable response-operator recovery

A simple explanation for CGC was that the model failed to recognize the held-out context. That explanation did not hold. We evaluated a contemporary zero-shot State model^6^ on five official held-context cytokine splits comprising 90 cytokines and 2,000 compact response features. On conventional response metrics, State substantially improved over a perturbation-mean baseline: *Δ*-Pearson increased from 0.259 to 0.489, while *Δ*-MSE changed only modestly from 0.0248 to 0.0244 (Fig. 2a).

**Figure 2.**
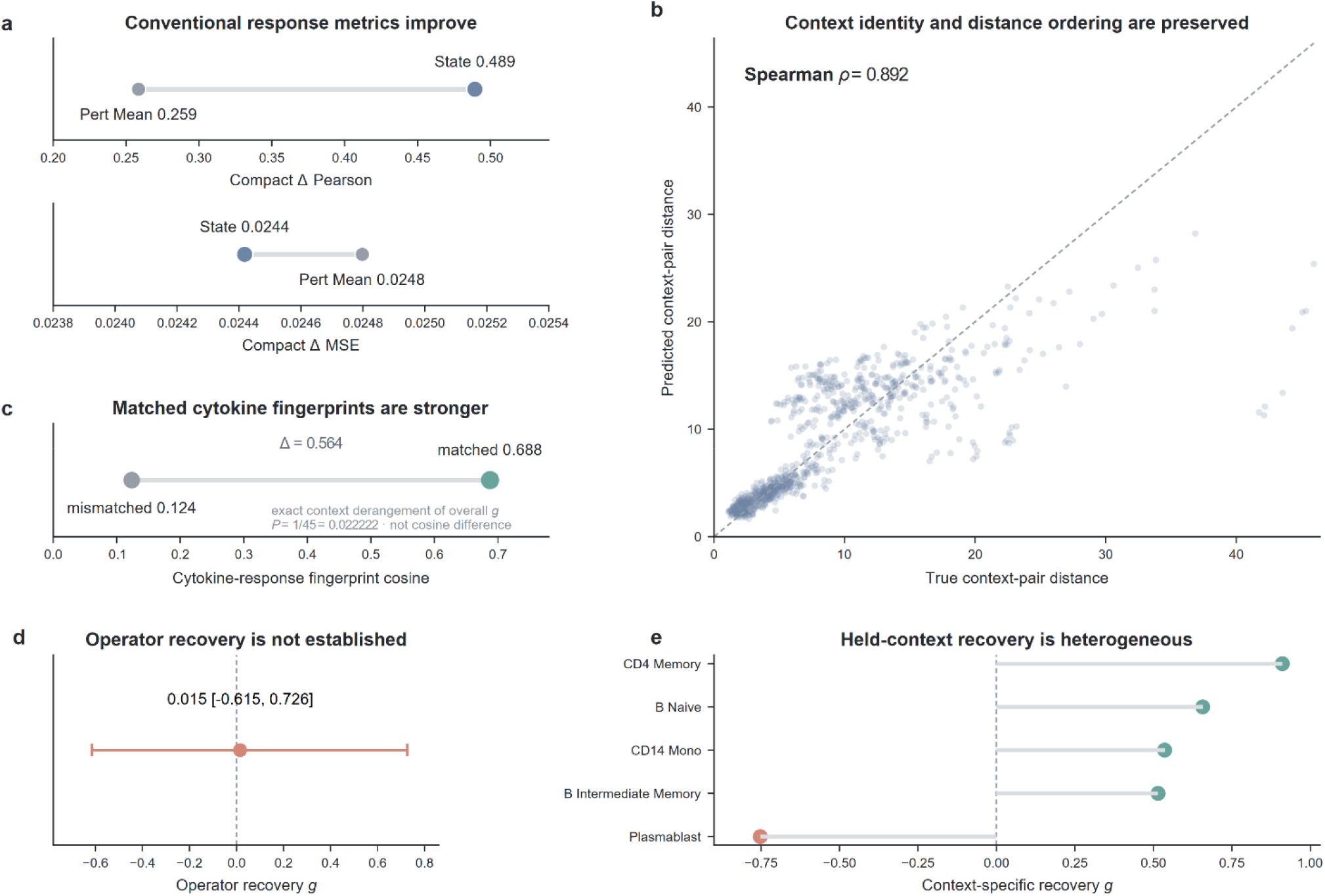
The identity–operator gap separates context identity from response-operator recovery. a, State versus Pert Mean on compact Δ-Pearson (0.489 versus 0.259) and Δ-MSE (0.0244 versus 0.0248). b, True versus predicted context-pair distances across 900 context-pair × cytokine observations (Spearman ρ=0.892). c, Within-context matched versus mismatched cytokine-response fingerprint cosine (0.688 versus 0.124); a separate exact context-derangement test of overall operator recovery gave P=1/45. d, Overall held-context operator recovery, g=0.0153 (prespecified cross-split uncertainty interval, −0.6146 to 0.7261), did not establish positive recovery. e, Per-context recovery; four contexts were positive, whereas Plasmablast was strongly negative (g=−0.753).

State also preserved the relational organization of contexts. Across 900 context-pair × cytokine comparisons^11^, predicted context distances closely tracked true distances (Spearman *ρ* = 0.892; Fig. 2b). Separately, within each held context, the cosine between matching predicted and true cytokine-response fingerprints was 0.688, compared with 0.124 for mismatched cytokines, a descriptive difference of 0.564 (Fig. 2c). An exact finite context-derangement test of overall operator recovery found zero exceedances among 44 unique derangements, giving the conservative add-one *P* = 1/45 = 0.022222.

Despite this strong preservation of context identity and ordering, positive novel-context operator recovery was not established. The prespecified recovery statistic was *g* = 0.0153 with a cross-split uncertainty interval from −0.615 to 0.726 (Fig. 2d). Four of five contexts had positive point estimates, but Plasmablast showed strongly negative recovery (*g* −0.753; Fig. 2e). Thus, State could recover substantial operator structure in selected contexts, but context identity did not guarantee reliable recovery. This favorable pattern did not replicate in an independent official zero-shot drug-response setting: only one of five Tahoe contexts showed positive recovery and overall recovery was −0.497 (hierarchical-bootstrap interval, −1.334 to −0.084; Extended Data Fig. 1a,b). State retained direction-sensitive signal, but context-specific operator-energy inflation degraded calibrated operator recovery (Extended Data Fig. 1c). We next tested whether denser empirical support could improve context-specific operator recovery.

A separate generic context-calibration analysis revealed a broad positive context tendency that failed intervention-, sentinel- and reference-context correspondence controls and showed negative bidirectional transfer (Supplementary Fig. 2a–d). Context identity and global context tendency were therefore insufficient to specify the intervention-conditioned response operator.

### Near-complete empirical support leaves the context-specific operator largely unresolved

If the identity–operator gap simply reflected sparse training coverage, then a sufficiently dense intervention × context matrix should make the remaining entries easy to infer. To test this directly, we constructed a strict prospective Experimental Compression analysis in the Tahoe-100M^12^ replicate core. The matrix contained 50 contexts and 93 exact drug-by-dose interventions (4,650 perturbation entries) measured on two independent plates. After fixed sample-local normalization and matched-control subtraction, each target response was sealed before episode-specific preprocessing, support and sentinel selection, hyperparameter selection, fitting and prediction. The model could use *m* complete reference contexts and *k* sentinel interventions measured in the target context, but the target response had no causal path to fitted quantities until final evaluation (Fig. 3a). Five adversarial integrity challenges—four hidden-target outcome replacements and one direct hidden-row-access challenge—left support selection, sentinel selection, hyperparameters and fitted predictions unchanged, establishing the prospective integrity of the pipeline (Extended Data Fig. 2a). Throughout the replicate-resolved Tahoe analyses, normalized recovery is 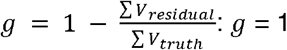 denotes complete recovery, *g* = 0 no gain over the matched reference, and *g* < 0 negative transfer. It is a cross-replicate signal-recovery statistic, not conventional R^2^.

**Figure 3.**
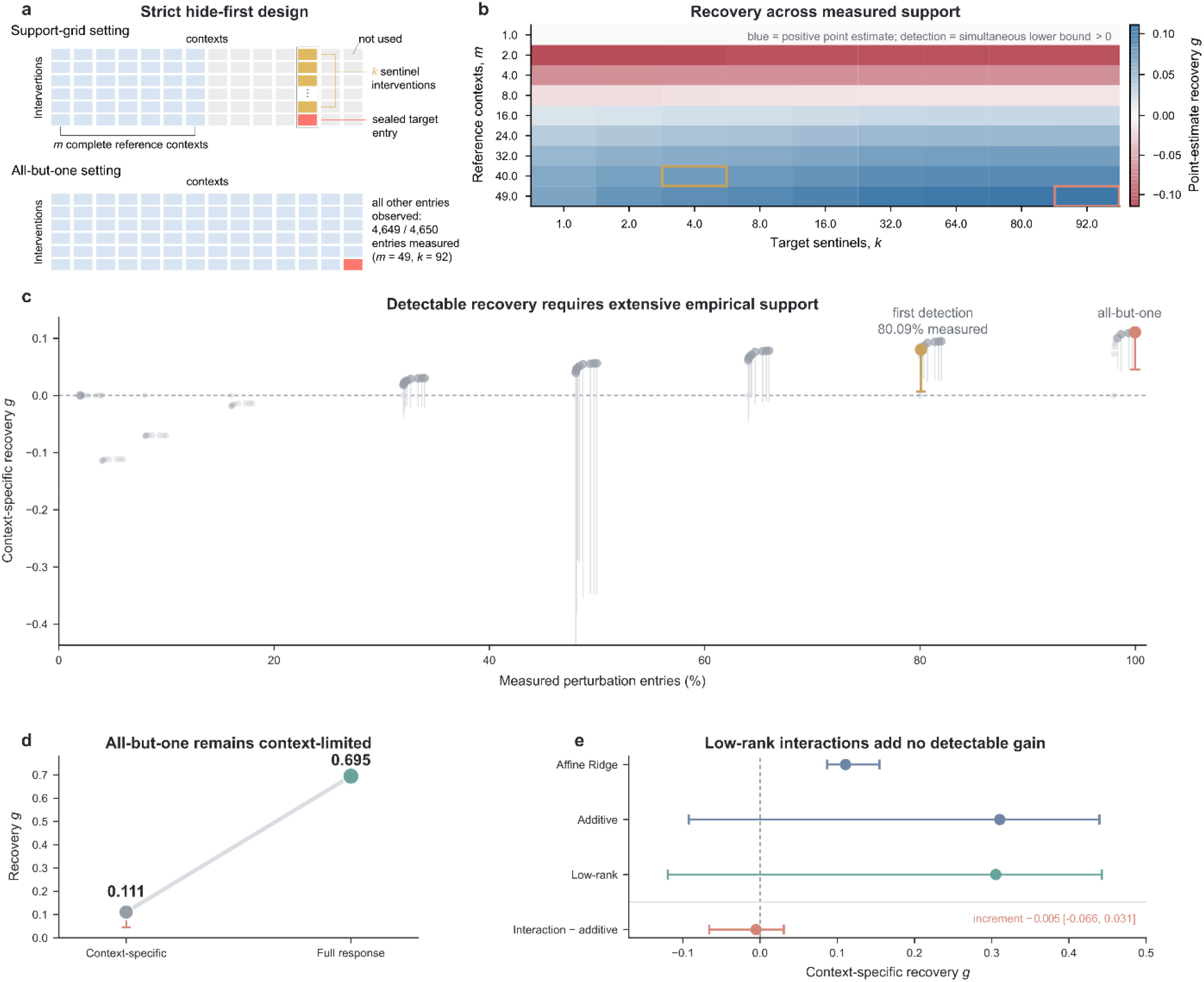
Near-complete experimental support leaves context-specific response structure largely unresolved. a, Strict hide-first Experimental Compression over 50 contexts × 93 interventions measured on two independent plates.Fixed sample-local normalization and matched-control subtraction preceded each episode; the target response was then sealed before episode-specific preprocessing, support and sentinel selection, hyperparameter selection and fitting; the inset shows the all-but-one limit, repeated for all 4,650 entries. b, Exact affine-Ridge recovery surface across the prespecified reference-context (m) × target-sentinel (k) support grid. The k=0 baseline was retained in the 90-point inferential family but omitted from the heatmap because recovery was identically zero. Outlines mark first detection (m=40, k=4) and all-but-one (m=49, k=92). c, Context-specific recovery versus measured-entry fraction; vertical segments show one-sided simultaneous 95% FWER lower bounds. Detectable recovery first appears after 3,724/4,650 entries (80.09%; g=0.0803; lower bound, 0.00688), and the 25%, 50%, 80% and 95% recovery thresholds are not reached. d, At all-but-one, full-response recovery is 0.6945, whereas context-specific recovery is 0.1107 (simultaneous lower bound, 0.04552). e, Under matched 99% target-hidden out-of-fold support, affine, additive and low-rank interaction models yield g=0.1107, 0.3104 and 0.3053. The interaction-minus-additive increment is −0.0051 (95% hierarchical-bootstrap interval, −0.0657 to 0.0308).

Across the preregistered 90-point *m* × *k* grid, recovery increased only slowly with measured support (Fig. 3b,c). The first point whose one-sided simultaneous 95% family-wise-error-rate lower confidence bound exceeded zero occurred at *m* = 40 and *k* = 4, corresponding to 3,724 of 4,650 measured perturbation entries (80.09% of the matrix). Even there, context-specific recovery was only *g* = 0.0803, with a simultaneous lower confidence bound of 0.00688. Thresholds for 25%, 50%, 80% and 95% recovery were not reached within the prespecified grid (Fig. 3c). Matched positive controls established sensitivity to shared and synthetic low-rank structure, and all four correspondence-null families were exceeded at the first-detection and all-but-one budgets (Extended Data Fig. 2b–d).

We then evaluated the strongest possible empirical-support limit of this design. In the all-but-one setting, each matrix entry was separately hidden while the remaining 4,649 of 4,650 entries were observed (Fig. 3a,d). Full-response recovery reached 0.6945, indicating that the shared and coarse perturbational response was highly reconstructable. By contrast, context-specific recovery remained 0.1107, with a one-sided simultaneous 95% FWER lower confidence bound of 0.0455 (Fig. 3d). Almost complete empirical coverage could therefore reconstruct much of what an intervention generally does while leaving most of the final context-specific deviation unresolved.

To test whether this bottleneck reflected estimator capacity, we repeated near-complete target-hidden completion with additive main effects and an explicit rank ≤32 context × intervention interaction under matched 99.0% out-of-fold support. Additive effects captured additional broad structure, but the explicit low-rank interaction term contributed essentially no incremental recovery beyond them (*Δ*g=−0.0051; 95% hierarchical-bootstrap interval, −0.0657 to 0.0308; Fig. 3e). Final fits used the full selected-rank range, contextwise interaction increments remained heterogeneous, and the same implementation recovered a matched rank-4 interaction almost completely (Extended Data Fig. 3a–c). Thus, dense support resolved broad additive tendencies without revealing the remaining pair-specific operator as a simple reproducible low-rank interaction.

Same-context anchors improved recovery only with broad reference-context support. Once broad reference-context coverage had established a useful response basis, a few anchors recovered a compact transferable component, yet context-specific recovery remained below 0.10 even under dense anchoring (Extended Data Fig. 4a–d). If neither near-complete matrix coverage nor direct measurements in the target context closed the gap, the next question was no longer simply how much data had been seen, but what counterfactual information the conditioning state actually specified.

### Baseline RNA contains counterfactual information, but at unequal resolution

We therefore asked whether baseline RNA specifies the full individual-specific response or only a coarser counterfactual structure. We tested this in the lymphoblastoid cell-line study of Lea et al.^13^, in which stable 1000 Genomes LCLs were exposed in parallel to dexamethasone (DEX) and ethanol control. After filtering, 342 biological LCLs had paired ETOH–DEX profiles across 10,157 processed expression rows representing 10,120 unique GeneIDs. The available public matrix was post-SVA voom-residualized; this experiment therefore tests the operational sufficiency of the available RNA representation under strict biological-LCL-disjoint prediction (Fig. 4a).

**Figure 4.**
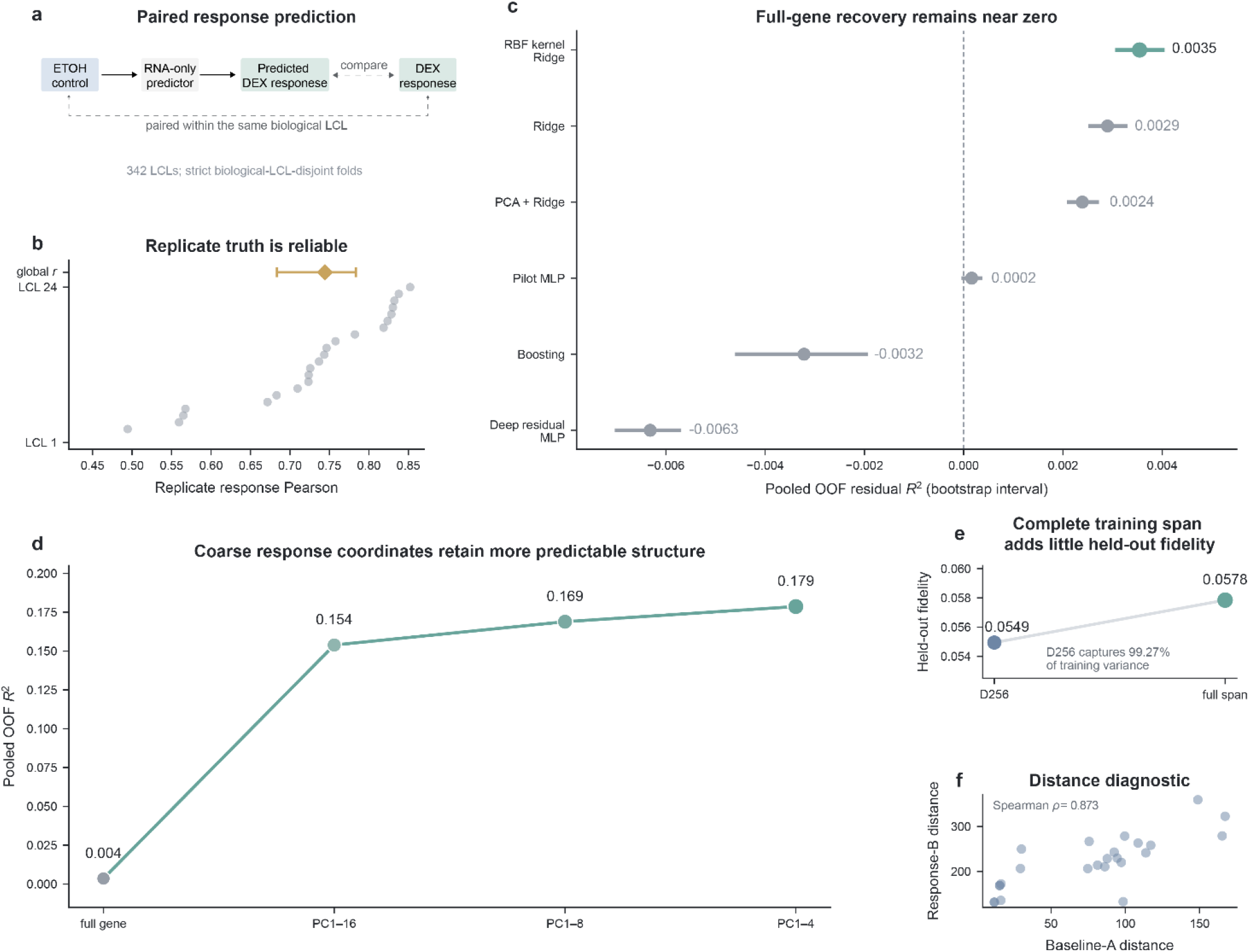
Baseline transcriptomic state supports coarse response prediction but yields little recovery of full individual-specific gene responses. a, Paired ETOH–DEX design in 342 biological LCLs with strict LCL-disjoint prediction. b, Measurement-disjoint replicate reliability in 24 replicate-complete LCLs; flattened residual-response correlation is 0.744 (95% LCL-bootstrap interval, 0.684–0.783). c, Full-gene out-of-fold residual R^2^ across six model families; the best tested model, RBF kernel Ridge, reaches 0.00354. d, Projecting the same frozen RBF predictions onto coarser response coordinates increases recovery: PC1–16, 0.1539; PC1–8, 0.1689; and PC1–4, 0.1787. e, A D = 256 training basis explains 99.27% of training variance yet retains only 0.054945 held-out fidelity; the complete training span reaches 0.057844. f, Pairwise baseline and independently measured response distances were descriptively correlated (Spearman ρ=0.873). Separately, baseline-nearest-neighbor pairs had smaller median independent-response distance than random comparisons (lower-tail empirical P=1/10,001).

Individual-specific response was reproducible. In 24 replicate-complete LCLs, the global flattened residual-response correlation between measurement-disjoint replicate sets was 0.744 (95% LCL-bootstrap interval, 0.684–0.783; Fig. 4b). Thus, the same DEX intervention elicited reproducible Individual-specific response was reproducible. In 24 replicate-complete LCLs, the global flattened residual-response correlation between measurement-disjoint replicate sets was 0.744 (95% LCL-bootstrap interval, 0.684–0.783; Fig. 4b). Thus, the same DEX intervention elicited reproducible individual-specific response differences beyond the shared treatment effect. We then trained a deliberately broad but prespecified model battery to predict the individual-specific DEX residual response from ETOH baseline expression. Full-gene out-of-fold *R*^2^ remained close to zero across Ridge (0.00290), PCA+Ridge (0.00239), MLP (0.00016), RBF kernel Ridge (0.00354), boosting (−0.00322) and a deep residual MLP (−0.00632; Fig. 4c). The best tested model therefore explained less than 0.4% of the full individual-specific gene-response variance.

A matched semi-synthetic sensitivity control produced monotonic, directionally correct gains as deliberately encoded baseline-dependent response structure was strengthened (Extended Data Fig. 5a). The control was used to establish directional sensitivity of the frozen pipeline rather than as a quantitative bound on the information content of the biological response.

The same frozen gene-level predictions achieved substantially higher recovery scores at lower response resolution. Using fold-local response principal components fit only within each outer-training fold, RBF kernel Ridge achieved pooled out-of-fold *R*^2^ values of 0.1787 for PC1–4, 0.1689 for PC1–8 and 0.1539 for PC1–16 (Fig. 4d). In a measurement-disjoint diagnostic, pairwise baseline and independent response distances were descriptively correlated (Spearman *ρ=*0.873); separately, baseline-nearest-neighbor pairs had smaller median independent-response distance than random comparisons (lower-tail empirical *P*=1/10,001 Fig. 4f). The gene-to-program hierarchy also persisted in an independently reprocessed non-SVA ARCHS4^15^ representation on the same 340 LCLs and the same prespecified outer folds (best full-gene recovery, 0.0108; PC1–4 recovery, 0.1475; Extended Data Fig. 5b). Full-gene recovery also remained near zero in AFR- and EUR-restricted analyses (Extended Data Fig. 5c). Baseline RNA retained coarse response-relevant information, although the tested RNA-only models recovered little of the full individual-specific gene response.

Importantly, low-dimensional response recovery did not imply high-fidelity reconstruction of the complete response. A 256-dimensional training-response basis explained 99.3% of training variance but retained only 5.5% of held-out individual-specific response; extending to the complete training-response span increased fidelity only to 5.8% (Fig. 4e; Extended Data Fig. 5d,e). The fine-scale individual response was therefore not concentrated in a transferable shared linear span.

### The pooled fine-scale residual response is globally high-dimensional

In the Lea analysis, baseline RNA supported coarse response prediction, yet the full gene-level displacement was nearly unrecovered. One remaining possibility was that the missing component was merely a compact latent correction that current models had failed to extract. We tested this in the full Tahoe replicate core—50 contexts, 93 shared interventions, 25,695 primary genes and two independent plates—where a reproducible context-specific operator was confirmed across multiple prespecified gene universes (Fig. 5a). We characterized residual-response structure in this complete, transductively defined matrix.

**Figure 5.**
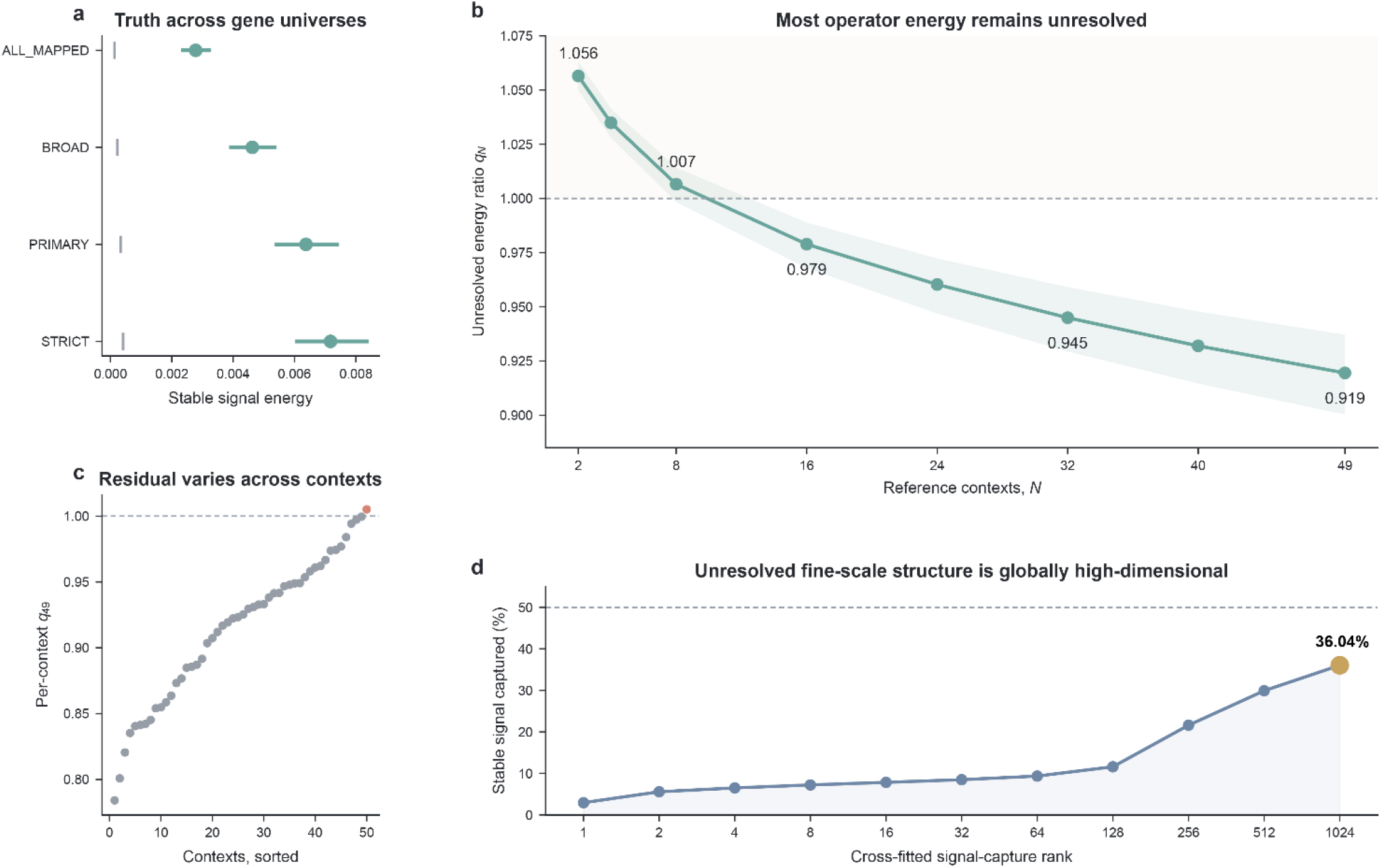
The pooled residual-response matrix remains globally high-dimensional at transcriptome scale. a, Replicate-stable operator truth across four prespecified gene universes: all mapped genes, the broad QC set, the primary gene set and the strict QC set. Points show observed signal energy and grey ticks the corresponding 95th-percentile shuffle values (empirical P = 1/5001 for each truth gate). b, Unresolved energy ratio q□ after transductive reconstruction from N = 2 to 49 reference contexts. At N = 49, q□ □ = 0.9195 (95% bootstrap interval, 0.9003–0.9371); q = 1 denotes no recovery and q > 1 negative transfer. c, Per-context maximal-support q□ □ across all 50 contexts, ranging from 0.7841 to 1.0052. d, Cross-fitted cumulative signal capture in the pooled residual-response matrix, with context–intervention pairs as rows and genes as columns, through rank 1,024. Rank 1,024 captures 36.04% of signal, and the 50%, 80% and 90% capture thresholds all exceed 1,024. Bases were fitted separately by plate and intervention fold; capture was scored by cross-plate agreement on held-out interventions.

We quantified unresolved structure with *q*_*N*_, the normalized residual-to-reference operator ratio. *q*_*N*_ =1 denotes no net recovery relative to the reference estimator, *q*_*N*_ <1 positive recovery and negative transfer; *q*_*N*_ < 1 is therefore not constrained to [0,1]. The pooled ratio decreased from *q*_2_ =1.0565 to *q*_8_ =1.0065 *q*_16_ =0.97899 *q*_32_ =0.94850 and *q*_49_ =0.9195 (95% bootstrap interval for q□ □, 0.9003– 0.9371; Fig. 5b). Even with 49 reference contexts, the corresponding recovery *g*_49_=1− *q*_49_ was only 0.0805. Residual energy persisted across all 50 contexts (Fig. 5c). Thus, broad contextual support reduced but did not eliminate the unresolved operator.

The pooled residual-response matrix was not compact. Cross-fitted low-rank reconstruction captured only 36.04% of the residual signal at rank 1,024, the maximum tested rank, and the ranks required to capture 50%, 80% or 90% of the signal all exceeded 1,024 (Fig. 5d). The remaining response variation was therefore globally high-dimensional across contexts and perturbations.

We also tested whether pharmacological labels explained the unresolved response structure or mechanism-aligned baseline modalities improved its prediction. The unresolved operator did not robustly collapse onto broad action, mechanism of action, target family or exact target (Supplementary Fig. 3a–c). None of the tested mechanism-aligned baseline modalities established incremental recovery beyond matched RNA-only prediction, although alignment often avoided degradation from generic feature concatenation (Supplementary Fig. 4a–d).

### Lowering output resolution increases counterfactual recovery

We next asked whether the demanded output resolution exceeded what the available conditioning information could support. Without retraining the Virtual Cell, we evaluated the same fixed Experimental Compression predictions either as 25,695-gene vectors or after deterministic projection onto five externally defined signed PROGENy pathway coordinates^16^ (MAPK, PI3K, JAK–STAT, p53 and NFκB). At the all-but-one budget, full-gene recovery was 0.110681 and pathway-coordinate recovery was 0.272814; the paired gain was 0.162133 (max-T simultaneous 95% interval, 0.074411–0.249855; Fig. 6a and Extended Data Fig. 6a). Because the prediction was held fixed, the gain identifies a resolution effect rather than added model capacity or input information.

**Figure 6.**
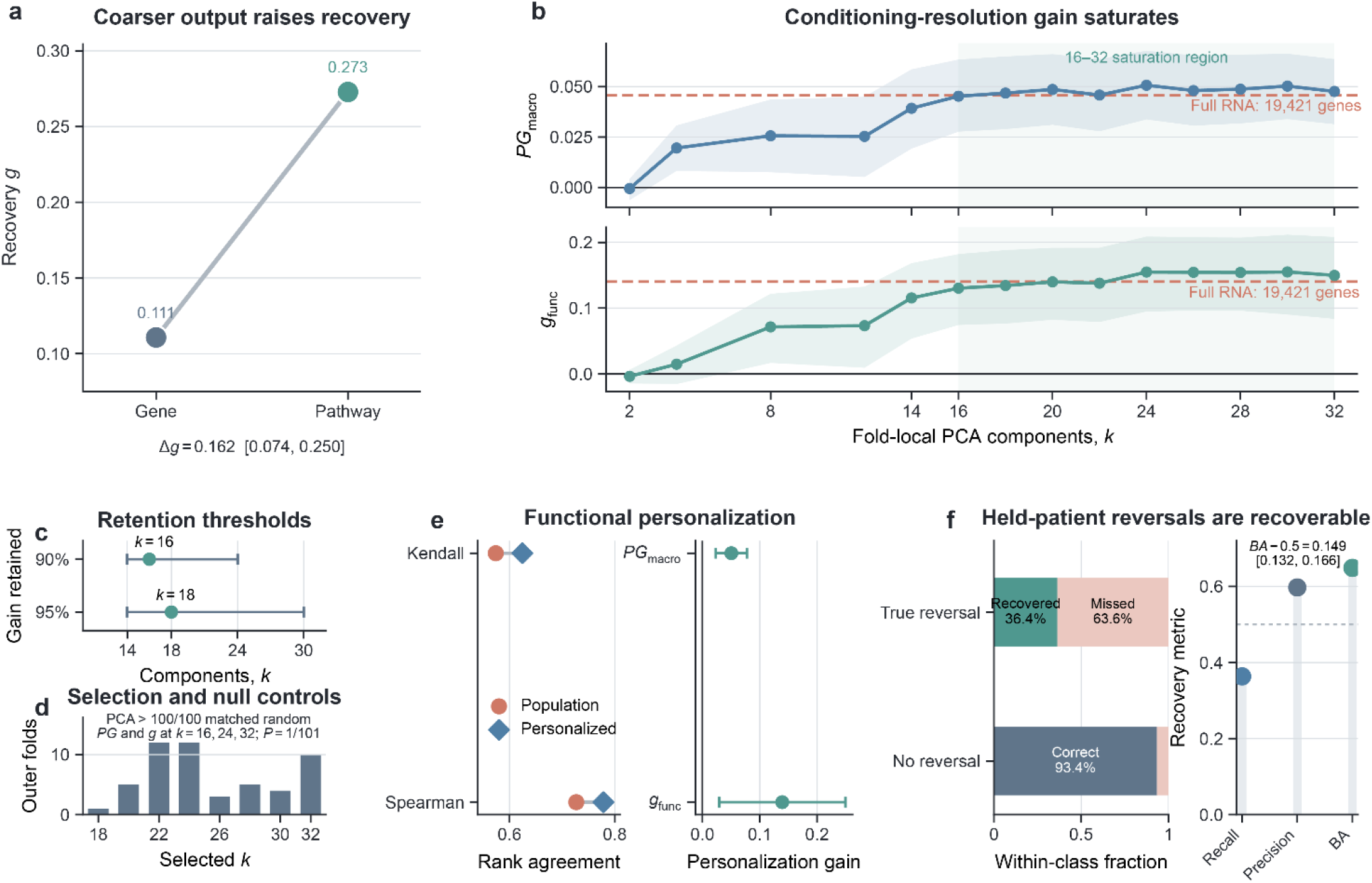
Counterfactual prediction is bounded by recoverable information resolution. a, The same fixed Experimental Compression predictions were evaluated at full-gene resolution or after projection onto five predefined signed PROGENy pathways. Recovery increased from 0.110681 to 0.272814 (paired gain, 0.162133; simultaneous 95% interval, 0.074411–0.249855). b, Patient-disjoint conditioning-resolution curves across fold-local PCA representations of 2–32 dimensions in 52 patients. Dashed lines denote reoptimized full-RNA performance and the shaded region marks the 16–32-dimensional saturation range. c, Smallest representations retaining 90% and 95% of both full-RNA endpoints were 16 and 18 components, respectively; horizontal segments show 90% bootstrap ranges. d, Inner-only model selection chose 18–32 components across all 52 outer folds; PCA16, PCA24 and PCA32 outperformed dimension-matched random projections for both endpoints. e, Baseline RNA improved within-patient drug ranking, with mean personalization gain *PG*_*macro*_=0.050423 and functional recovery *g*_*func*_ =0.139289. f, Among 3,045 held-patient drug-pair reversals, 36.4% were recovered; 93.4% of non-reversals were correctly retained (balanced accuracy,0.648844). All drug outcomes are ex vivo organoid responses.

Across the complete 14-pathway PROGENy vocabulary, the full span represented 3.43% of the reproducible gene-level operator, compared with 2.97% for the predefined five-pathway panel. Exhaustive evaluation of all 16,383 non-empty pathway subsets showed that pathway count explained 43.1% of fidelity variation but only 0.4% of recoverability variation; the predefined five-pathway panel lay at the 98.6th fidelity percentile but the 32.6th recovery percentile among five-pathway subsets (Extended Data Fig. 6b–f). Recoverability was therefore composition-dependent rather than a generic consequence of dimensionality reduction.

We next examined a distinct held-context scaling task, withholding all target-context responses and predicting them from baseline state and *N*_ref_ response-observed reference contexts, rather than the entrywise completion used in Fig. 3. At maximal observed support (*N*_ref_ = 49), gene- and pathway-level recovery reached 0.168976 and 0.361825, respectively (Extended Data Fig. 7a–c). The selected continuing-power gene fit projected approximately 62,811 reference contexts, each measured for all 93 interventions, to reach *g*=0.5. The selected pathway fit approached a ceiling of 0.416142; a matched continuing-power sensitivity fit projected *g*=0.5 at approximately 414 reference contexts (Extended Data Fig. 7d). Lower output resolution improved observed recovery, while extrapolated thresholds depended on model form.

### Predictive information saturates below nominal measurement resolution

We next tested the complementary conditioning-side prediction of the resolution principle in the metastatic colorectal-cancer organoid cohort^14^. Baseline RNA was measured across 19,421 genes, but measurement granularity need not equal the scale at which information about future response is transferable. Using the same 52 patients, 24 drugs, strict leave-one-patient-out folds and interaction-capable multi-output Ridge model as the application below, we replaced the full baseline profile with fold-local principal-component representations derived from full baseline RNA and spanning 2–32 dimensions. Full-RNA prediction was reoptimized under the same nested training-side procedure, so the experiment varied representation resolution while holding the outcome, estimator family, population reference and held-patient information set fixed (Fig. 6b–d and Extended Data Fig. 9).

Predictive performance rose rapidly with the first components and then entered a broad 16–32-dimensional plateau. Personalization gain increased from −0.000560 at two components to 0.045239 at 16 components, whereas *g*_*func*_ increased from −0.003731 to 0.130360. No adjacent step within 16–32 reached the prespecified practical effect scales of 0.010 for personalization gain or 0.030 for *g*_*func*_ . Full RNA achieved 0.045672 and 0.140286, respectively, and did not add a material increment over PCA representations of 16, 20, 24, 28 or 32 dimensions (Fig. 6b and Extended Data Fig. 9a–e).

Operational thresholds made this mismatch explicit. The smallest representation reaching at least 90% of both full-RNA endpoints was 16 components, and the corresponding 95% threshold was 18. Patient-level bootstrap distributions placed the median joint thresholds at 16 and 18, with 90% ranges of 14–24 and 14–30, respectively (Fig. 6c). Ratios of the nominal gene-coordinate count to these representation dimensions were 1,214 and 1,079, respectively. Training-side model selection independently chose 18– 32 components in all 52 outer folds, and PCA representations at 16, 24 and 32 dimensions each exceeded all 100 dimension-matched random projections for both endpoints (empirical *P* = 1/101; Fig. 6d and Extended Data Fig. 9f). Predictive performance therefore saturated in a compact representation derived from the full transcriptome.

Saturation did not imply that any coarse representation was sufficient. Fourteen PROGENy pathway activities exceeded matched random pathway vocabularies but retained substantially less personalization than full RNA; adding 16–32 residual transcriptomic components produced material gains (Supplementary Fig. 5). The operational information scale was therefore much more compressible than the measured transcriptome, yet richer than a small canonical pathway vocabulary.

### Baseline molecular state supports partial patient-specific drug prioritization

We next asked whether the information recoverable from baseline state supported a functional, patient-specific decision. The Kryeziu metastatic colorectal-cancer organoid resource^14^ contains 213 organoids from 102 patients and 24 harmonized drugs with complete author-reported sensitivity scores; the primary RNA-seq intersection comprised 91 organoids from 52 patients. Every preprocessing, tuning and prediction operation used strict leave-one-patient-out separation, and organoids from the same patient never crossed folds. The endpoint was ex vivo drug sensitivity, not clinical response.

The population drug ordering already achieved mean within-patient Kendall τ_b_ = 0.5740. Baseline-RNA prediction increased this to 0.6244 and improved ranking in 37 of 52 patients. Mean personalization gain was 0.050423 (simultaneous 95% interval, 0.023299–0.077547; max-T-adjusted *P* = 0.00030), and patient-specific functional response recovery was *g*_*func*_ = 0.139289 (simultaneous 95% interval, 0.029092–0.249487; adjusted *P* = 0.01160; Fig. 6e). Baseline molecular state therefore supplied reproducible but incomplete information beyond the population drug response.

Among 14,320 held-patient drug-pair comparisons, 3,045 (21.264%) reversed the training-population ordering. The RNA model recovered these reversals with recall 0.364, precision 0.597, balanced accuracy 0.649 and Matthews correlation coefficient 0.363; balanced-accuracy gain above chance was 0.148844 (simultaneous 95% interval, 0.131745–0.165944; adjusted *P* = 0.00010; Fig. 6f). Top-k summaries, near-tie behavior, correspondence nulls and patient-level heterogeneity are reported in Extended Data Fig. 8.

Directionally consistent personalization was reproduced in the secondary HTA2.0 cohort of 118 organoids from 50 patients (mean gain, 0.060534; simultaneous 95% interval, 0.034257–0.086810; adjusted *P* = 0.00010; Extended Data Fig. 8e). Patient-specific functional prediction remained useful despite limited recovery of fine-scale molecular responses across the preceding analyses.

## Discussion

Our study establishes recoverable biological resolution as a criterion for evaluating virtual-cell predictions. Measurement resolution, the resolution at which predictive information is recoverable and the resolution requested of a model need not coincide. CGC exposes this distinction: a model can preserve cellular identity and shared perturbation effects while losing the response differences that distinguish biological contexts. The resulting gap is not an all-or-none failure of prediction. Useful response structure can remain recoverable at coarser resolutions.

The most direct evidence comes from changing the resolution of the same predictions. In the drug-response matrix, predefined pathway summaries achieved higher normalized recovery than gene-level profiles without retraining the predictor. This gain reflects which response features were requested, not additional input information or model capacity. Pathway identity mattered more than pathway count, indicating that a coarser output must still capture the relevant biology. The gene-to-program hierarchy in lymphoblastoid cells, reproduced after independent RNA reprocessing, further showed that resolution dependence was not confined to predefined pathways or a single biological system. By contrast, measuring 4,649 of 4,650 entries left context-specific recovery at 0.1107 in the affine completion analysis. Dense coverage did not make fine-scale responses reliably recoverable in this setting. The input-side result was complementary: 16 and 18 transcriptomic components retained 90% and 95% of full-RNA drug-response prediction performance across both organoid endpoints. Together, these results show why experimental coverage, measurement dimensionality and recoverable biological detail cannot substitute for one another.

Classical conditional prediction provides a theoretical interpretation of these findings. Under squared-error loss, the optimal predictor is the conditional mean of the response given the available information^17^. When that information leaves context-specific response variation unresolved, averaging over possible responses can contract their amplitude and geometry towards a shared rule. Empirical anchors can improve recovery by constraining this variation, while selected pathway or program coordinates can remain predictable when much of the full response is unresolved. The relevant information may therefore support different levels of detail for different response components.

Recoverable resolution describes the level of biological detail that can be reproducibly predicted from the available information.

These conclusions are operational rather than information-theoretic limits on prediction from a complete transcriptome. Recoverability depends on the cohort, task, estimator and available information. The Lea hierarchy replicated after independent non-SVA reprocessing, but its semi-synthetic control remained sample- and model-limited. Low-rank and full-span linear analyses tested specific explanations without exhausting nonlinear mappings. Tahoe residual-response anatomy was transductive, whereas Experimental Compression tested prospective completion. Pathway summaries capture selected biological features rather than high-fidelity substitutes for the full transcriptome. Likewise, the compact organoid representations were derived from full RNA: their component counts describe predictive performance thresholds, not the size of a reduced assay or an intrinsic biological dimension. These distinctions matter when transferring the framework to a new system or prediction task.

A resolution-aware approach gives virtual-cell development a concrete objective: recover finer biological features with the same experimental support, or maintain recovery while reducing the measurements required. Representation learning may concentrate predictable response structure into useful biological coordinates. Decoding these representations into gene-expression profiles does not itself establish finer context-specific recovery. Benchmarks should therefore assess recovery at both the representation and output levels in unmeasured context–perturbation combinations. Experimental design should test which additional observations improve that recovery. Gene-level outputs remain appropriate where supported; coarser outputs are useful when they retain the biology needed for the task. Patient-specific drug ranking, for example, can help prioritize ex vivo testing even when detailed molecular responses remain unresolved. Matching prediction resolution to recoverable information makes it possible to evaluate experimental substitution in terms of the biological question being answered, rather than the dimensionality of the transcriptome being generated.

## Methods

### Conceptual definitions

For a biological context *c* and intervention *p*, we write the post-perturbation state as *Y*_*p,c*_=*X*_*c*_ + *Y*_*p,c*_ *Δ*_*p,c*_,where *X*_*c*_ is the measured baseline state and *Δ* _*p,c*_ is the perturbation response. We decompose the response into a shared component *Δ* _*p*_, estimated across contexts for the same intervention, and a context-specific component *Γ* _*p,c*_. The shared component is estimated using only the training or reference contexts allowed in each analysis. CGC refers to systematic contraction or loss of the reproducible geometry of *Γ* _*p,c*_ under held-context prediction.

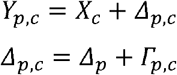

For a fixed context *c*, the full response operator is the map *p* ⟼ *Δ*_*p,c*_, and its context-specific component is the map *p* ⟼ *Γ*_*p,c*_. For a finite panel of *P* interventions and *G* response coordinates, the matrix of context-specific responses is:

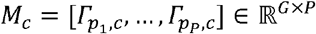

If *e*_*j*_ selects intervention *p*_*j*_, then 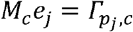. Columns index individual perturbations, without assuming additive effects of combined perturbations. Recovery metrics are evaluated on the response vectors or contrasts specified for each analysis.

Measurement resolution is the detail recorded by an assay; prediction resolution is the detail requested of a counterfactual prediction. Output-side recoverable resolution is the biological detail reproducibly predicted for a specified information set, representation, estimator and held-out evaluation. On the input side, we measure the representation size at which held-out performance saturates. Identifiable and predictive identifiability refer to empirically supported recovery under these conditions.

### Conditional-prediction interpretation

*S* denotes all information available at prediction time, including baseline state, intervention identity, reference contexts, empirical anchors and other permitted variables. Under squared-error loss, the Bayes-optimal predictor of the context-specific response component is its conditional expectation. The remaining component *U* is unresolved by that information. This decomposition interprets CGC; it is not a fitted model. For a lower-resolution projection *PΓ*, conditional uncertainty may be smaller than for the full response vector *Γ*, which can make programs or pathways more recoverable^17^.

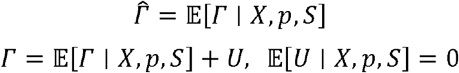

### Replicate-stable recovery

Where independent plates or response halves were available, we measured operator recovery using cross-replicate signal energy. For a truth response D and prediction 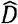, residuals 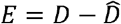 were formed independently in the two replicate directions. Reproducible truth and residual energies were estimated from cross-products between replicate representations. Recovery was defined as 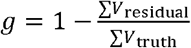. This quantity is not constrained to [0,1]: *g < 0* denotes negative transfer. Supplementary Table 3 lists the prespecified statistical units, resampling schemes and multiplicity families. This recovery statistic is distinct from conventional R^2^.

### Technical variation and batch handling

Response construction preserved matched controls and independent replicates. In the T-cell analysis, each source pseudobulk row was normalized as log1p(CPM), and each targeting-guide-by-run response was formed relative to the equal-weight NTC mean matched within donor, stimulation state and 10x run. In Tahoe, each plate was normalized and DMSO-subtracted separately; Plate 6 and Plate 14 remained separate throughout fitting and evaluation, and recovery was estimated from cross-plate signal. Scaling, PCA and other learned transformations were fitted on training units and applied unchanged to held-out units.

### Primary human CD4+ T-cell CGC analysis

We used author-provided donor × stimulation-state × 10x-run × guide pseudobulks from primary human CD4+ T-cell CRISPRi Perturb-seq, with four donors and three stimulation states (Rest, Stim8h and Stim48h). Prediction was at the pseudobulk level. The source inventory contained 12,730 unique perturbed gene identifiers; the guide-disjoint reproducibility set contained 3,186 targets and 38,232 response vectors per half. Model evaluation used a prespecified primary response set with 9,386 intervention targets and 9,082 response genes. Aggregation weighted targeting guides and guide-run rows equally. Guide-disjoint halves were used for truth estimation and nuisance nulls. The assay universe, including exact-zero filtering, was fixed from the complete processed inventory before leave-one-donor-out evaluation; context PCA, the response basis, hyperparameter selection and model fitting were outer-training-only.

Reproducible response structure was evaluated using replicate-disjoint donor partitions and stimulation contrasts. Ridge, bilinear and MLP models used four leave-one-donor-out folds. A rank-256 response basis was fitted on training responses only. Prediction used the held donor’s untreated/NTC state but no held-donor perturbation outcomes. Truth-aligned amplitude α and cosine were summarized by a prespecified 10,000-draw intervention bootstrap with all four donors fixed. Donor-specific response *R*^2^ (Fig. 1e) was one minus the model-to-shared-response squared-error ratio. Both errors were computed on perturbation-by-stimulation interaction tensors, double-centered across perturbations and stimulation states for each gene; the reference was the equally weighted mean of the three training donors. Squared errors were pooled across perturbations, states, genes and held donors before seed-level scores were averaged. For Fig. 1f, each predicted interaction tensor was projected onto the shared-response tensor, using inner products pooled across those same coordinates and donors. Shared and residual energies were the squared norms of the projection and its orthogonal residual. Seed-level component energies were averaged before dividing shared energy by their sum. Construction and held-donor information sets are summarized in Supplementary Fig. 1 and Supplementary Table 2.

For an outer fold holding out donor *d*, let *h*_*d,s*_ ∈ ℝ^*H*^, *H* ≤ 8, denote the held donor’s NTC state in stimulation state *s* after standardization and PCA fitted on the three training donors. Let *U* _*− d*_ ∈ ℝ^9082 × 256^ be the response basis fitted on training responses, and let *µ*_*− d*_,_,*p,s*_ be the mean response to intervention *p* in state *s* across those training donors. Each model predicts residual coefficients and adds the same intervention- and state-specific anchor:

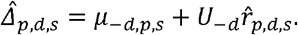

Ridge fits a separate coefficient matrix for each covered intervention:

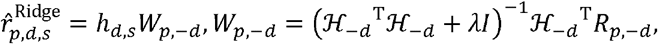

Here, ℋ_*− d*_ contains the nine training-donor-by-state NTC rows and *R*_*p, − d*_ contains the corresponding 256-dimensional anchored response residuals. The Ridge penalty was selected by leave-one-training-donor-out validation within each outer fold.

The bilinear residual head is

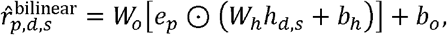

with a 32-dimensional learned intervention embedding and a 32-dimensional context embedding. The MLP residual head is

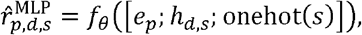

Here, *f*_*θ*_ has two 256-unit GELU layers and dropout 0.1. Both neural heads output 256 coefficients for the reconstruction above. The response basis was learned from centered training responses; coefficient projection follows the frozen direct projection convention without subtracting the saved PCA mean.

### State held-context evaluation

The State analysis used five official zero-shot held-context artifacts, 90 cytokines and 2,000 compact response features. Models were not refitted locally. State was compared with the matched perturbation-mean baseline on compact mean-vector *Δ*-Pearson and *Δ*-MSE. We measured Spearman correlation between true and predicted distances for ten unordered context pairs and 90 cytokines *(n =* 900) comparisons). Within each held context, a 90 × 90 predicted-by-true cytokine-response cosine matrix gave the matched (diagonal) and mismatched (off-diagonal mean) descriptive fingerprint scores. The finite context-correspondence test enumerated 44 unique context derangements; with zero exceedances, the conservative add-one P value is 1/45=0.0222. Without donor or replicate metadata, operator recovery used the prespecified deterministic cross-split uncertainty interval, not replicate-calibrated truth.

### State independent stability audit

We evaluated the same target-excluding recovery metric in official zero-shot ST-SE-Tahoe artifacts across five held contexts, without local refitting. Held contexts were the stability units; uncertainty used the prespecified hierarchical bootstrap. ST-HVG was the prespecified sensitivity representation. We checked axis correspondence, target exclusion and leakage invariance.

### Tahoe-100M replicate core

The Tahoe replicate core comprised 50 cell-line contexts, 93 exact shared drug-by-dose interventions, 25,695 primary genes and two independently normalized plate replicates (Plate6 and Plate14). The core contained 4,650 unique treatment entries and 9,500 pseudobulks derived from 13,390,662 retained cells; two DMSO wells per context and plate provided controls. Plate6 and Plate14 remained separate during fitting and were used as independent replicate directions for recovery metrics.

### Strict prospective Experimental Compression

Experimental Compression predicted hidden matrix entries. After fixed sample-local normalization and matched-control subtraction, each target response was hidden before episode-specific preprocessing, support and sentinel selection, hyperparameter selection, fitting and prediction. Support was defined by *m* complete reference contexts and *k* sentinel interventions measured in the target context. The structured budget was *B* = 93*m* + *k* over a 50 x 93 = 4,650-entry perturbation matrix. Prespecified values of were 1, 2, 4, 8, 16, 24, 32, 40 and 49; *k* values were 0, 1, 2, 4, 8, 16, 32, 64, 80 and 92, yielding 90 support points (Supplementary Table 5). The primary predictor was affine Ridge. Five integrity challenges—four target-outcome replacements and one direct hidden-row-access challenge— confirmed that hidden outcomes could not affect fit-side objects or predictions.

For target context *c*, intervention *p*, and plate *r* ∈ {6,14}, let *S*_*m*_ be the allowed set of *m* complete reference contexts and *K*_*c,p*_ the allowed target-context sentinel interventions. The estimator predicts the hidden response by an affine mixture of reference-context responses to the same intervention:

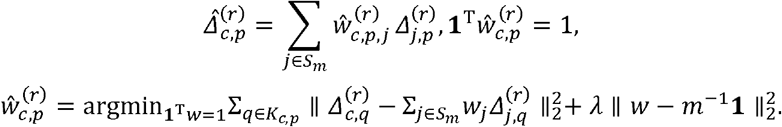

The target response 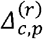 is absent from the sentinel set and from fitting. The estimator uses uniform reference-context weights when *k* = 0 or *m* = 1; otherwise *λ* is selected using only reference contexts in the episode’s observed support set. Plate 6 and Plate 14 are fitted and evaluated separately.

At each support point, recovery *g* was aggregated across the 4,650 separately hidden target episodes. The prespecified hierarchical bootstrap gave one-sided simultaneous 95% FWER lower confidence bounds over the 90-point family. Support thresholds for 25%, 50%, 80% and 95% recovery were obtained by inverting these bounds. The all-but-one design corresponds to *m* = 49, *k* = 92, for which 4,649 entries are observed and exactly one target entry remains hidden.

### Low-rank interaction completion audit

We tested model capacity with 100-fold entrywise prediction, retaining 99.0% of the 4,650 entries in each fit. The matched comparison included the frozen affine model, an additive context-plus-intervention model and a regularized CP interaction model with ranks 2, 4, 8, 16 and 32; rank and regularization were selected within each outer mask using only observed-entry inner masks. Each model predicted every entry out of fold on both plates; Plate6 and Plate14 remained separate for replicate-stable recovery. We checked target-outcome invariance, caches and convergence. A matched synthetic rank-4 interaction tested implementation sensitivity; uncertainty used the prespecified hierarchical bootstrap. The matched-affine comparator reused the frozen context- and plate-specific reference-only *λ* values selected in the earlier target-sealed analysis, then refitted its weights on each 99%-observed mask.

For the matched completion audit, let y denote the gene-level response indexed by context, intervention and gene. The additive model and the rank-R CP interaction extension were parameterized as

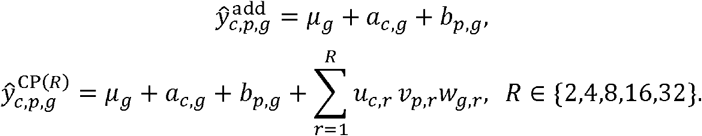

### Same-context anchoring analysis

This prespecified secondary analysis reused the target-hidden Experimental Compression episodes. Prespecified reference-context supports were *m* = 4, 16 and 40. Early anchor levels were *k* =0, 1, 2, 4 and 8, with the dense *k* = 92 endpoint testing extensive anchoring. Anchoring effects were paired changes in context-specific recovery within each stratum using the prespecified multiplicity-corrected inference procedure. At *m* = 40, the *k* = 2 and *k* = 8 gains were 0.076671 (simultaneous 95% FWER interval, −0.000561–0.153904) and 0.082467 (0.029907–0.135027), respectively; the first two anchors accounted for 93.0% of the estimated *k* = 8 gain. At *m* = 4, *k* = 8 produced a negative effect of −0.069603. Even at *k* = 92, recovery remained below 0.10 for all three reference-support levels (*m* = 4, −0.070003; *m* =16, 0.030226; *m =* 40, 0.094835).

### Lea paired-LCL RNA-only sufficiency analysis

The Lea analysis used author-processed post-SVA voom residual expression from paired ETOH and DEX samples. After filtering, 342 biological LCLs (177 AFR and 165 EUR) had paired profiles across 10,157 expression rows representing 10,120 unique GeneIDs. For each LCL *i*, the response was 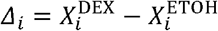. The outer-training mean DEX response was subtracted to define the individual-specific target *R*_*i*_. Models received only the ETOH baseline profile; genotype, ancestry and DEX-side predictors were excluded. All preprocessing and response bases were fitted inside biological-LCL-disjoint outer folds.

For outer fold f, the training-only shared DEX response and the held-LCL individual-specific residual target were defined by

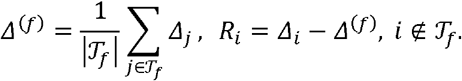

The prespecified model battery comprised Ridge, PCA+Ridge, MLP, RBF kernel Ridge, boosting and a deep residual MLP. Primary full-gene performance was pooled out-of-fold residual *R*^2^. Program analyses projected the same frozen full-gene RBF kernel-Ridge predictions and held-out responses onto PC1–4, PC1–8 and PC1–16 coordinates of a response basis fitted inside each outer-training fold. A separate set of 24 two-to-two replicate-complete LCLs was used for measurement-disjoint truth and distance diagnostics.

### ARCHS4 non-SVA robustness analysis

We repeated the Lea analysis using ARCHS4 v2.5 Kallisto-derived counts from independent reprocessing of the original reads, without SVA. Both representations used the same 340 identity-resolved LCLs, prespecified outer folds and 10,110 matched GeneIDs. ARCHS4 pseudocounts were normalized per sample and aggregated outcome-blind across multiple GEO versions within each line-by-treatment group before model fitting; the residual target, fold-local response-program bases and evaluation pipeline were otherwise unchanged. The original post-SVA representation was rerun on the same 340-LCL core, and all ten leakage-invariance checks passed. ARCHS4 is not an exact reconstruction of the Lea STAR/HTSeq pre-SVA matrix.

### Lea training-response span analysis

In each outer fold, we fitted a thin singular-value decomposition of training individual-specific responses and projected held-out responses onto the first 256 components or the full non-zero training span. Numerical ranks were 270–272. We verified full-rank training reconstruction, orthogonal projection identities and agreement with the frozen oracle-fidelity implementation. Held-out truth was used only to score projections. This tests transfer of the training-derived linear span, not nonlinear representability.

### Lea semi-synthetic sensitivity control

The semi-synthetic control retained the Lea baseline matrix x and the same biological-LCL-disjoint cross-validation and model pipeline, but replaced the response with a target containing a baseline-dependent latent component. Response reliability was calibrated near the empirical replicate benchmark of 0.7443, and indexed the strength of encoded baseline dependence. Sensitivity was assessed from out-of-fold R^2^, endpoint gain and monotonicity across . For RBF kernel Ridge, R^2^ increased from −0.003676 at *λ* = 0 to 0.036710 at *λ* = 1; for Ridge, R^2^ increased from −0.001240 to 0.025912. Both models had positive endpoint gains in all 10 comparisons and Spearmant (*λ*, R^2^) = 1. No tested model reached the preregistered positive-control threshold of R^2^ ≥ 0.25. This tests pipeline sensitivity, not the information content of the biological response.

### Tahoe operator reconstruction and dimensionality

Tahoe operator anatomy used the prespecified full-transcriptome operator with a transductive target definition. For *N* reference contexts, *q*_*N*_ was defined as the pooled normalized unresolved-operator ratio:

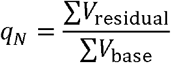

*q*_*N*_ = 1 denotes no net recovery relative to the reference estimator, *q*_*N*_ < 1 positive recovery and *q*_*N*_ > 1 negative transfer. We additionally report g_N_ = 1 − *q*_*N*_ as an equivalent recovery measure.

After maximal-support affine reconstruction, we pooled residuals into a matrix of context–intervention pairs by 25,695 genes. Responses were centered across interventions within each training or evaluation subset. Across five intervention folds, basis-fit matrices contained 3,350–4,150 rows. A gene-space basis was fitted on one plate and training interventions, then evaluated using cross-products of residual projections from both plates on held-out interventions, with plate roles exchanged. Captured and total energies were summed across folds and plate directions before taking their ratio. Rank refers to this pooled matrix rather than the within-context operator matrix *M*_*c*_. Cross-fitted reconstruction was evaluated over ranks 1–1,024. When 50%, 80% or 90% capture was not reached by rank 1,024, the corresponding threshold was reported as >1,024.

### Mechanistic attribution of the unresolved operator

We tested biological groupings of the prespecified unresolved Tahoe operator. Cross-plate analyses preserved replicates and tested broad pharmacological action, mechanism of action, target family, exact target and a prespecified set of 424 genotype–drug relationships. Independent clusters were counted at each level. Exact-target evidence had only three independent clusters and was prespecified as power limited. Program projections, including PROGENy, were secondary confirmation analyses. The primary output was the highest hierarchy level with robust cross-plate support; none met the prespecified criterion.

### Mechanism-aligned multimodal prediction

We tested whether intervention-aligned non-RNA features improved prediction of the prespecified residual target beyond RNA. Panels included wiring, pathway/activity, exact-target phosphoprotein, exact-target total protein, a functional dependency oracle and a joint multimodal representation (Supplementary Table 6). Each panel compared RNA, generic multimodal and mechanism-aligned features on its own complete-case set of contexts and interventions. Missing modalities were not zero-imputed in primary analyses. Estimators were prespecified Ridge and bilinear models. Paired recovery gains were measured against the matched RNA-only comparator. Fold-local synthetic controls estimated modality-specific sensitivity; analyses were omitted where no valid fold-local power implementation was available. Non-positive results in power-limited panels do not establish absence of the corresponding biology.

### Counterfactual Resolution analysis

We reused Experimental Compression predictions without retraining and verified full-gene recovery before projection. Five externally defined signed PROGENy pathways—MAPK, PI3K, JAK–STAT, p53 and NF_κ_B—were mapped to the 25,695-gene primary axis and L2-normalized. Each hidden target was evaluated in gene space and after projection onto the five pathways. Primary analysis used the all-but-one budget (m=49, k=92); m=40, k=4 was a prespecified secondary budget. Both resolutions used the same replicate-stable estimator. Each replicate was projected separately before computing cross-replicate truth and residual energies; replicates were not averaged into a denoised truth.

Let W denote the G-by-K matrix whose columns are the L2-normalized signed pathway weight vectors. Truth and prediction used the same fixed projection,

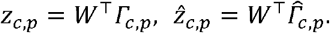

The replicate-stable recovery statistic g was then evaluated unchanged in the projected coordinate space.

The paired pathway-minus-gene contrast was evaluated over the same 4,650 held-out target episodes and corrected with the prespecified four-member maximum-absolute max-T simultaneous 95% confidence interval. This compares gene and pathway recovery under fixed predictions, not pathways against arbitrary low-dimensional projections.

### Pathway fidelity and resolution analysis

Pathway fidelity used the frozen 14-pathway PROGENy weight matrix on the 25,695-gene axis (Supplementary Table 4). Each subset defined an orthogonal projector onto its signed pathway span, using a fixed singular-value-decomposition pseudoinverse tolerance. Plate6 and Plate14 truth representations were projected separately, and pathway fidelity was defined as the fraction of the full cross-plate reproducible operator signal retained in that span. All 16,383 non-empty subsets were enumerated for fidelity; recoverability was defined for the 16,379 subsets with positive projected truth denominators. We verified the primary five-pathway panel and used the frozen replicate-stable estimator, leaving recovery undefined for non-positive projected truth denominators. Pathway-count summaries pooled all compositions of each size.

For a pathway subset S, the orthogonal projector onto its signed-weight span and the corresponding cross-plate biological fidelity were

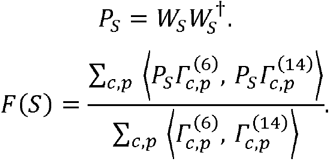

Here † denotes the Moore-Penrose pseudoinverse evaluated with the fixed singular-value-decomposition tolerance used in the frozen analysis.

Held-context scaling withheld all target-context responses; prediction used its baseline and responses from *N*_ref_ reference contexts. The RBF kernel-ridge predictor used DMSO profiles centered across genes and scaled to unit norm, with prediction weights adjusted to sum to one. For *N*_*ref ≥ 8* ridge_ penalties (0.001, 0.01, 0.1, 1, 10) and kernel-scale factors (0.25, 0.5, 1, 2, 4) were selected by fivefold validation within reference contexts, minimizing cross-plate prediction-error energy. Kernel scale was the factor divided by the median positive squared distance between reference profiles; smaller support sets used penalty 0.1 and factor 1. The same predictions were evaluated in gene space and the prespecified five pathways. We fitted *q* = 1 − *g* across observed support levels by least squares, comparing constant (*q* = *q*_0_) continuing-power 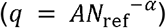 finite-floor power 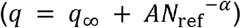 and saturating exponential 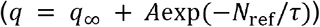 functions. The selected fit minimized leave-one-support-level-out RMSE, with the corrected Akaike information criterion as tie-breaker; continuing power was selected for genes and the saturating exponential for pathways. The continuing-power sensitivity fitted that form at both resolutions, independently of the selected pathway fit. For Extended Data Figure 7 curves and contrasts, the hierarchical bootstrap resampled target contexts, then interventions within each context.

### Patient-specific ex vivo drug-prioritization analysis

The application used the Kryeziu et al. metastatic colorectal-cancer patient-derived organoid resource (Mendeley Data v3, DOI 10.17632/hr94h42xdc.3; GEO GSE294511). We used processed expression/count data and author-reported drug-sensitivity scores, not raw sequencing. Before modeling, an outcome-blind availability check selected 24 drugs with complete harmonized sensitivity measurements. The primary RNA-seq cohort contained 91 organoids from 52 patients; the secondary HTA2.0 cohort contained 118 organoids from 50 patients. Eligible organoids with expression and complete drug screens were averaged within patient.^14^

Prediction used leave-one-patient-out splits; preprocessing, filtering, dimensionality reduction, tuning and fitting used only training patients. For held patient i, the 24-drug population reference was the mean response vector among training patients. Multi-output Ridge fitted a separate baseline-expression coefficient vector for each drug to predict residual responses relative to this reference, allowing patient-specific drug ordering. The primary application used full baseline RNA, with PCA20 plus Ridge retained as a prespecified sensitivity analysis.

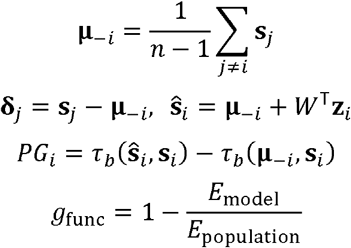

Personalization gain was averaged across held patients. Functional recovery used patient-balanced squared error relative to the training-patient population reference. Reversal classification considered drug pairs whose held-patient ordering opposed the training-population ordering and reported recall, precision, balanced accuracy and Matthews correlation coefficient. Patient-level bootstrap uncertainty was computed from the frozen out-of-fold prediction table; the three-endpoint primary sign-flip family used maximum-absolute simultaneous correction. Two within-fold training-patient correspondence nulls separately shuffled baseline RNA profiles or complete residual-response profiles. Both preserved drug labels and within-patient response-profile coherence. Null add-one probabilities are per-metric; primary sign-flip inference is simultaneous across three endpoints.

### Conditioning-resolution saturation analysis

Conditioning analysis reused the primary application’s 52-patient RNA-seq cohort, 24-drug matrix, patient grouping and outer folds. The baseline matrix contained 19,421 genes. Within each outer fold, genes were standardized using training patients only. Principal-component analysis was refitted within every inner-training split and on the complete outer-training set. Fixed dimensions were 2, 4, 8, 12, 14, 16, 18, 20, 22, 24, 26, 28, 30 and 32; the full standardized RNA matrix was the nominal gene-level comparator. Patient-grouped inner cross-validation selected the Ridge penalty and, for the dense selected model, component count.

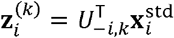

Outer-training loadings projected the standardized held-patient baseline without refitting. All representations used the same population-residual target and multi-output Ridge estimator. Saturation denotes flattening of held-patient performance for this task and estimator. Component counts describe full-RNA representations, not intrinsic transcriptome dimensions or reduced assay sizes.

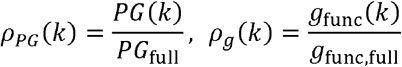

Retention was relative to reoptimized full-RNA performance for each endpoint. The joint operational threshold at q = 0.90 or 0.95 was the smallest component count at which both retention ratios reached or exceeded q. Patient-level bootstrap resampling estimated threshold distributions. Early and late gains compared 2–16 with 16–32 components; adjacent late gains were evaluated against prespecified practical scales of 0.010 for personalization gain and 0.030 for functional recovery. Inner-only selected dimensions were recorded in every outer fold. At 16, 24 and 32 dimensions, 100 outcome-blind orthonormal random projections provided dimension-matched controls.

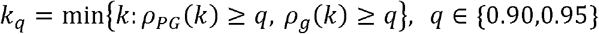

We also projected the training-standardized baseline into the rank-14 span of the full PROGENy vocabulary (17,349 of 19,421 genes mapped). PROGENy14 was compared with full RNA, PCA14, 100 14-dimensional random projections and 100 matched random pathway vocabularies. Fine residual components orthogonal to the PROGENy span were then added at 0, 4, 8, 16 and 32 dimensions. All projections and residual bases were fitted training-side only.

### Statistical analysis and validation policy

Split units and statistical units were prespecified. Bootstrap and permutation schemes are summarized in Supplementary Table 3. For finite permutation spaces, the number of unique permutations, not Monte Carlo draws, bounded P values. For Experimental Compression, inference used one-sided simultaneous 95% FWER lower confidence bounds over the 90-point family.

## Code availability

CGC-authored analysis and reporting code for all manuscript-facing analyses are available at https://github.com/yongqih/cgc-counterfactual-resolution and Zenodo: **10.5281/zenodo.22664234** under release tag v1.0.1. The deposited release includes prespecified split definitions, final model and evaluation configurations, figure source-data assembly, statistical resampling and null procedures, and Supplementary Table generation. Scientific entrypoints are classified as full analyses, reconstruction from frozen predictions, official-artifact summaries or table generators; source-table integrity checks are listed separately. CGC-authored code and documentation are provided under the MIT License. Third-party datasets, model artifacts and resource-derived materials retain their upstream licenses and access terms, as documented in the accompanying notices.

## Data availability

All biological datasets and official prediction artifacts analyzed in this study are publicly accessible, and no private outcome data are required. The primary human CD4+ T-cell Perturb-seq data are available through the CZI Virtual Cells Platform and under GEO GSE314342, SRA SRP643211 and BioProject PRJNA1359008 (https://doi.org/10.64898/2025.12.23.696273). The cytokine data underlying the State analysis are available as arcinstitute/State-Parse-Filtered, and the exact official compact evaluation artifacts are available as arcinstitute/ST-SE-Parse and arcinstitute/ST-HVG-Parse. The independent State–Tahoe evaluation artifacts are available as arcinstitute/ST-SE-Tahoe and arcinstitute/ST-HVG-Tahoe, with the linked arcinstitute/State-Tahoe-Filtered dataset. These State resources are used through the exact public representations and versions listed in Supplementary Table 1 and remain subject to their upstream non-commercial and acceptable-use terms.

Tahoe-100M is available as tahoebio/Tahoe-100M and through the Arc Virtual Cell Atlas (https://doi.org/10.1101/2025.02.20.639398). The analyses used the public plate-resolved pseudobulk representation and the filtered raw-count Plate 6 and Plate 14 objects, as specified in Supplementary Table 1. The paired lymphoblastoid cell-line study is available under GEO GSE207049 and SRA SRP383837; the exact author-processed matrix used here is archived at Zenodo record 6595427 (https://doi.org/10.5281/zenodo.6595427). The independent RNA reprocessing used the ARCHS4 v2.5 human gene-count matrix (human_gene_v2.5.h5; https://doi.org/10.1038/s41467-018-03751-6).

Fixed pathway definitions were obtained from PROGENy v1.17.3 (https://doi.org/10.1038/s41467-017-02391-6). Mechanism-aligned features used Reactome release 97 (https://doi.org/10.5281/zenodo.21383214), HGNC RRID:SCR_002827, DepMap Public 26Q1 hotspot and damaging mutation matrices, the 26Q1 Chronos gene-effect matrix, MCLP RPPA500 and the harmonized Sanger 2022 mass-spectrometry proteomics resource. Exact public identifiers, versions, representations and access terms for every resource are provided in Supplementary Table 1 and the accompanying dataset-accession manifest.

The metastatic colorectal-cancer organoid pharmacogenomic resource is available from Mendeley Data v3 (https://doi.org/10.17632/hr94h42xdc.3) and GEO GSE294511. The analysis used public processed expression or count matrices and author-reported drug-sensitivity tables; no raw sequencing files are redistributed.

Source-data tables for Main Figures 1–6, Extended Data Figures 1–9 and Supplementary Figures 1–5 are supplied with the article and on GitHub and Zenodo: **10.5281/zenodo.22664304** under release tag v1.0.1. Third-party data and model artifacts are linked rather than relicensed; the CGC-authored release code is licensed separately under MIT.

## Author contributions

Y.H. conceived the study, designed and implemented the computational analyses, performed the statistical analyses, generated the figures and wrote the original manuscript. H.W. contributed to methodological discussion and manuscript modification. C.L. contributed clinical expertise, helped develop clinically grounded application scenarios, and advised on the clinical interpretation and translational implications of the study. P.W. supervised the study, provided scientific guidance, contributed to interpretation of the results and revised the manuscript. All authors reviewed and approved the final manuscript.

## AI-assisted tools

Generative AI tools (OpenAI ChatGPT and Codex) were used to assist with code drafting and debugging, methodological brainstorming and language editing. All analytical decisions, statistical evaluations, interpretation of results and final manuscript content were reviewed and verified by the authors.

## Extended Data figure legends

**Extended Data Fig. 1.**
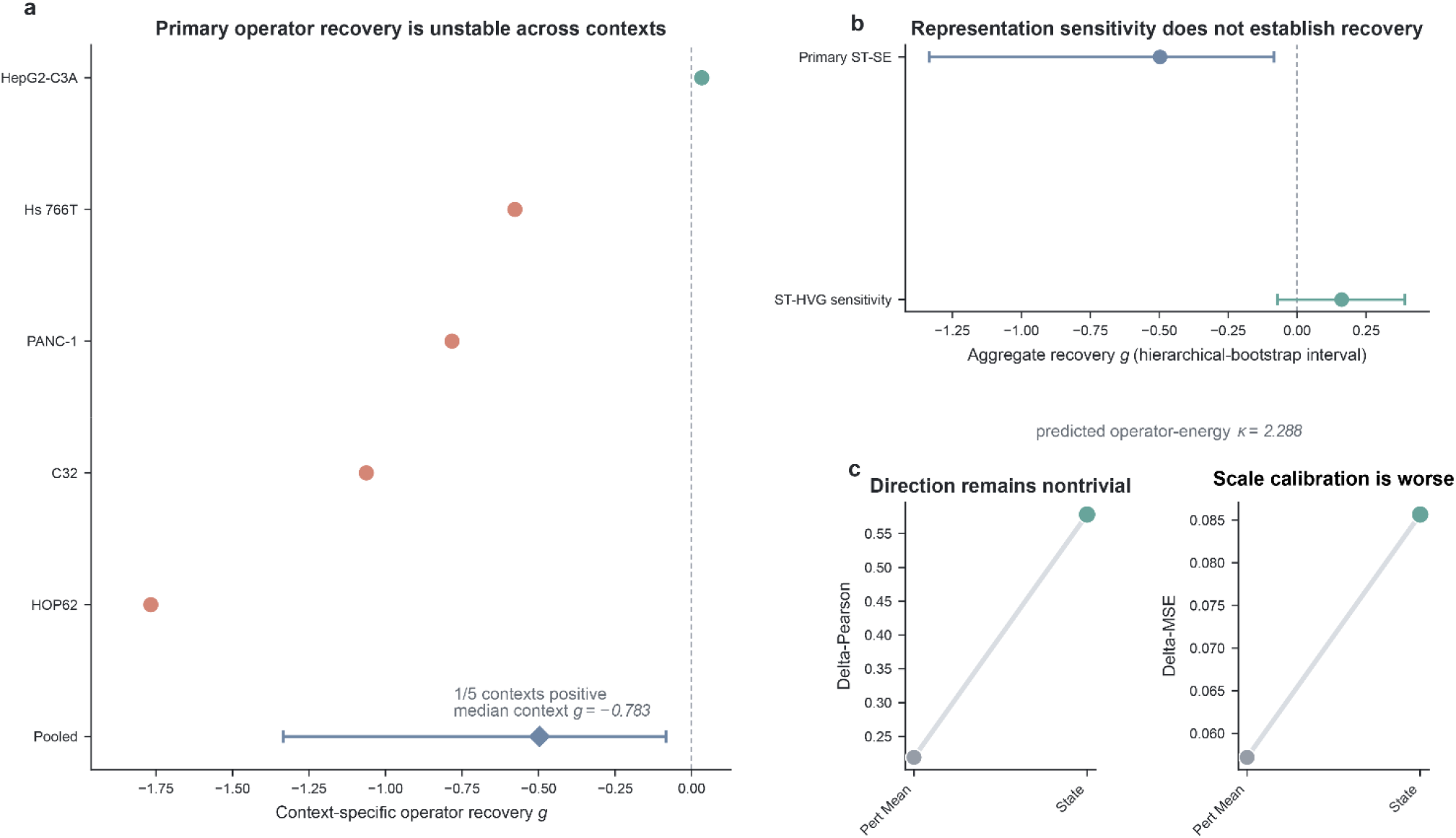
State retains response direction but not stable operator recovery in an independent drug-response setting. a, Context-specific operator recovery for five Tahoe held contexts under the primary ST-SE representation; one of five point estimates is positive and pooled recovery is g=−0.4968 (95% hierarchical-bootstrap interval, −1.3342 to −0.0836; median context g=−0.7828). b, Aggregate recovery for primary ST-SE and the prespecified ST-HVG sensitivity (g=0.1617; interval, −0.0713 to 0.3914), which does not establish positive recovery. c, State improves Δ-Pearson over Pert Mean (0.578 versus 0.219) but worsens Δ-MSE (0.0856 versus 0.0572); predicted context-specific operator energy is inflated relative to true operator energy (κ=2.288).

**Extended Data Fig. 2.**
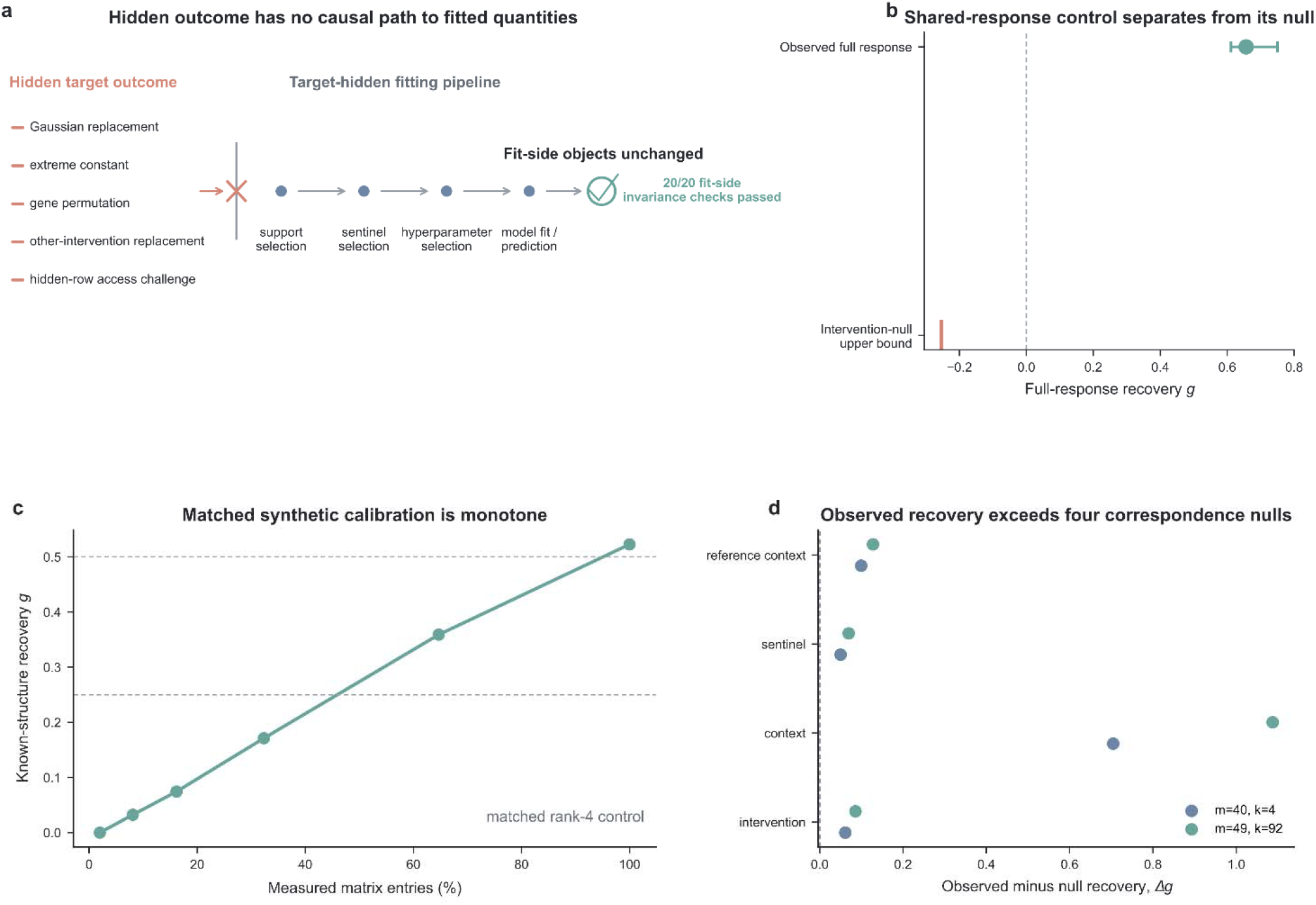
Experimental Compression is outcome-sealed and sensitive. a, After fixed sample-local normalization and matched-control subtraction, four hidden-target outcome replacements plus a direct hidden-row-access challenge left fit-side objects and predictions unchanged (20/20 checks). b, At m=49, k=0, the shared-response control gives full-response g=0.6565 (95% bootstrap interval, 0.6113–0.7502); the intervention-null upper bound is −0.2541. c, The support-grid rank-4 synthetic control reaches g=0.5227 at all-but-one. d, Recovery exceeds four correspondence-null families at m=40, k=4 and m=49, k=92.

**Extended Data Fig. 3.**
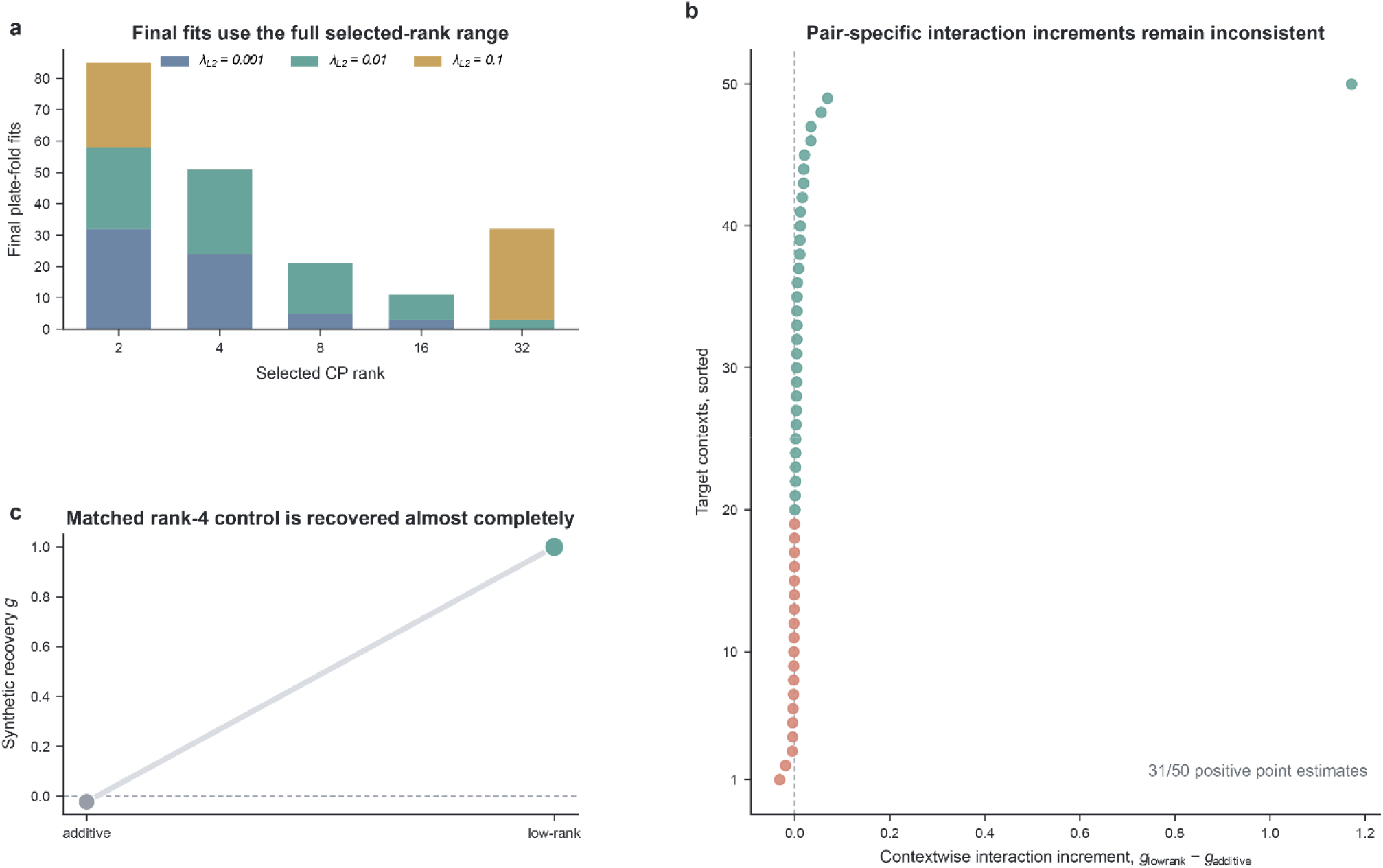
Active low-rank capacity does not recover the pair-specific interaction. a, Selected CP ranks and L2 penalties across 200 final plate-fold fits span the full prespecified rank grid (2–32). b, Contextwise interaction increments g_low-rank − g_additive across 50 contexts; 31 of 50 point estimates are positive, but effects remain heterogeneous. c, Under the distinct 99%-observed low-rank implementation control, a matched rank-4 synthetic interaction is recovered almost completely (low-rank g = 0.9998; gain over additive, 1.0217). All 200 final refits converged.

**Extended Data Fig. 4.**
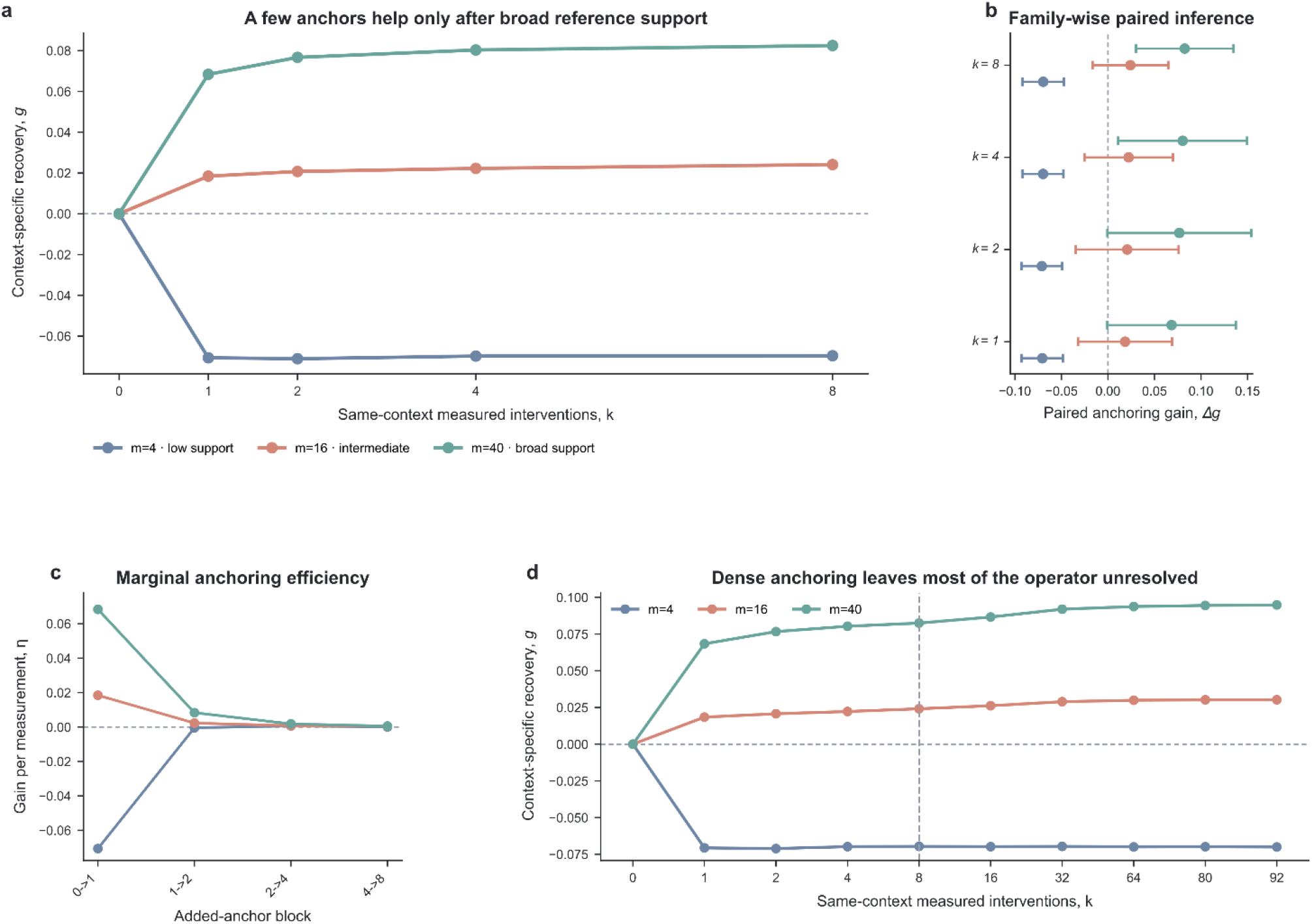
Same-context anchoring requires broad reference support. a, Recovery for k = 0, 1, 2, 4 and 8 target-context anchors at m = 4, 16 and 40. Anchors harm recovery at m = 4 but help at m = 16 and 40. b, Paired gains with simultaneous 95% FWER intervals; at m = 40, gains at k = 2 and k = 8 are 0.0767 and 0.0825. c, Marginal gain per anchor block is front-loaded under broad support. d, Through k = 92, recovery remains below 0.10 at every reference-support level.

**Extended Data Fig. 5.**
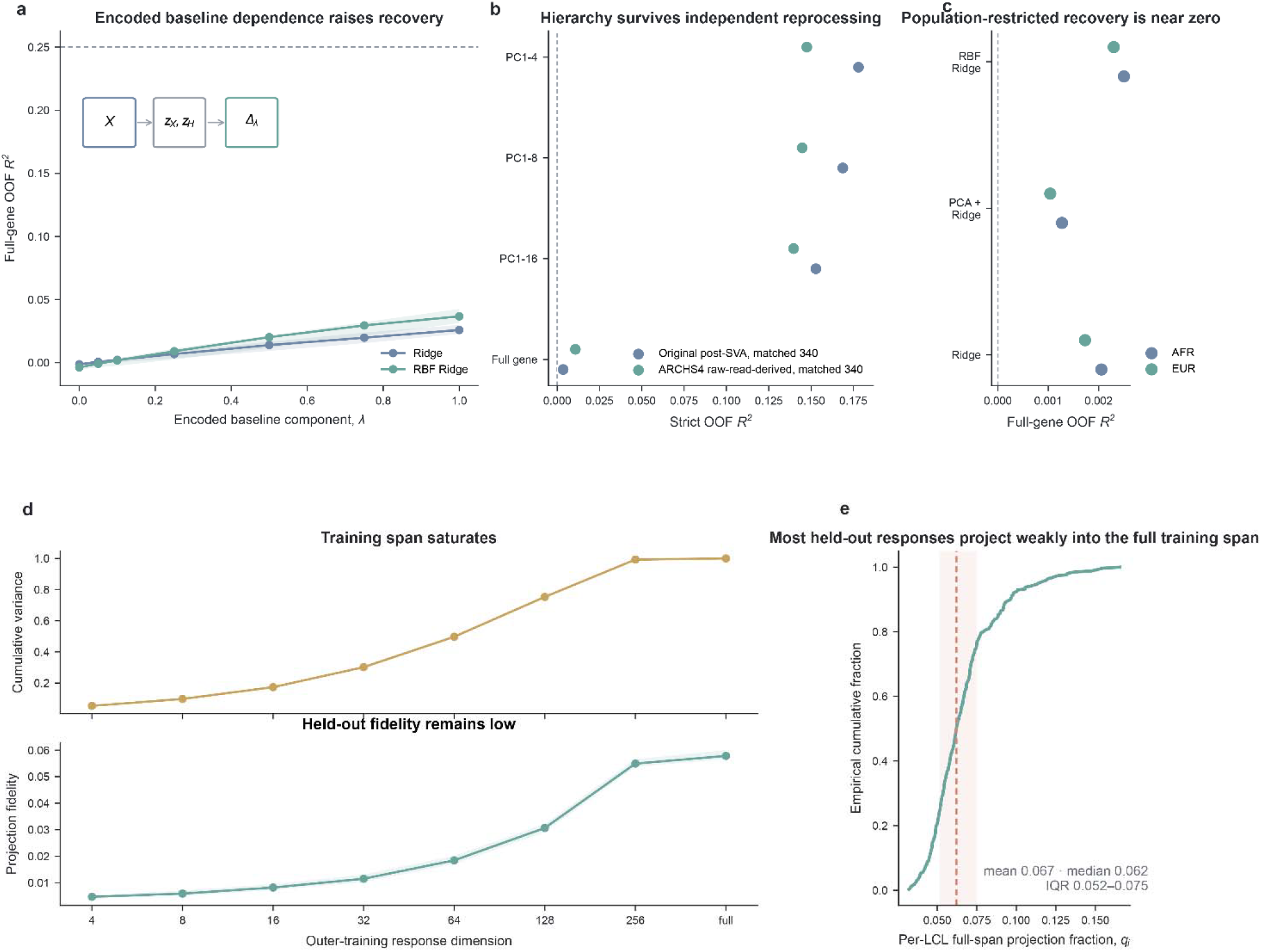
The Lea resolution hierarchy survives sensitivity, reprocessing, population and span analyses. a, Medians and interquartile ranges across ten prespecified seeds in matched semi-synthetic data. Full-gene out-of-fold R^2^ increases monotonically with encoded baseline dependence for Ridge and RBF Ridge but remains below the preregistered 0.25 threshold. b, The gene-to-coarse hierarchy persists in matched post-SVA and independently reprocessed ARCHS4 profiles from 340 LCLs. c, Full-gene out-of-fold R^2^ remains near zero in AFR- and EUR-restricted analyses. d, A training-response basis that captures 99.27% of training variance at D = 256 yields held-out fidelity of 0.0549; the complete numerical span reaches 0.0578. e, Per-LCL full-span projection fractions remain low (mean, 0.067; median, 0.062; interquartile range, 0.052–0.075).

**Extended Data Fig. 6.**
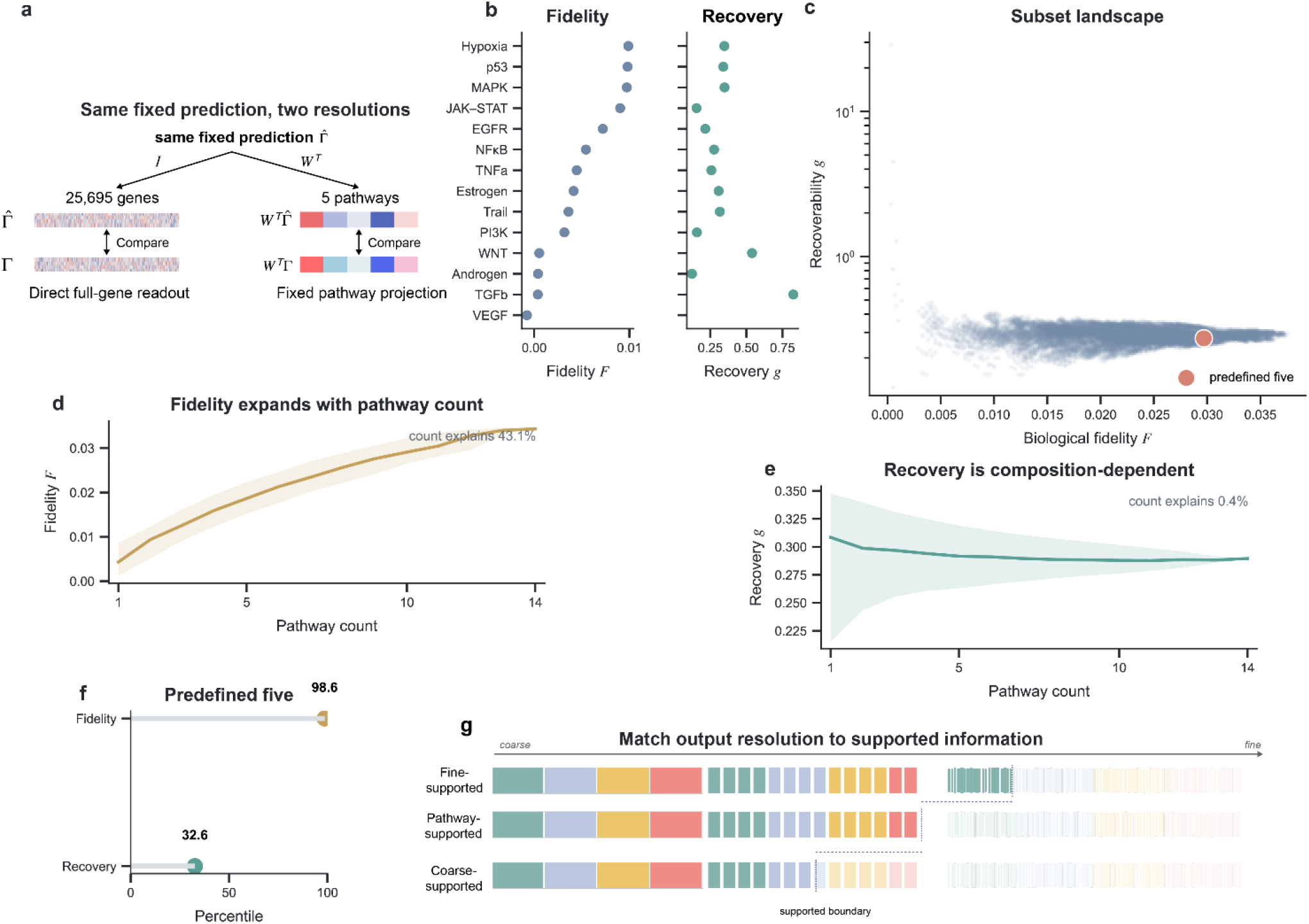
Composition organizes pathway fidelity and recoverability. a, Conceptual schematic of resolution matching under the same fixed prediction. b, Descriptive fidelity and recoverability across the 14 individual PROGENy pathways; VEGF recovery is undefined because its cross-plate projected-truth denominator is non-positive. c, Full landscape of 16,379 pathway subsets with positive projected-truth denominators. d,e, Pathway-count summaries for fidelity and recovery; fidelity rises and saturates with count, whereas recovery remains composition-dependent. Shading shows the descriptive interquartile range (25th–75th percentiles) across exhaustively enumerated pathway subsets at each pathway count. f, The predefined five-pathway panel is at the 98.6th fidelity percentile and 32.6th recovery percentile among 2,002 five-pathway subsets. g, Counterfactual output should retain gene-level detail for fine-supported components and use pathway-level output when only coarse structure is supported. Fidelity includes all 16,383 non-empty subsets; four non-positive denominators are omitted only from recovery.

**Extended Data Fig. 7.**
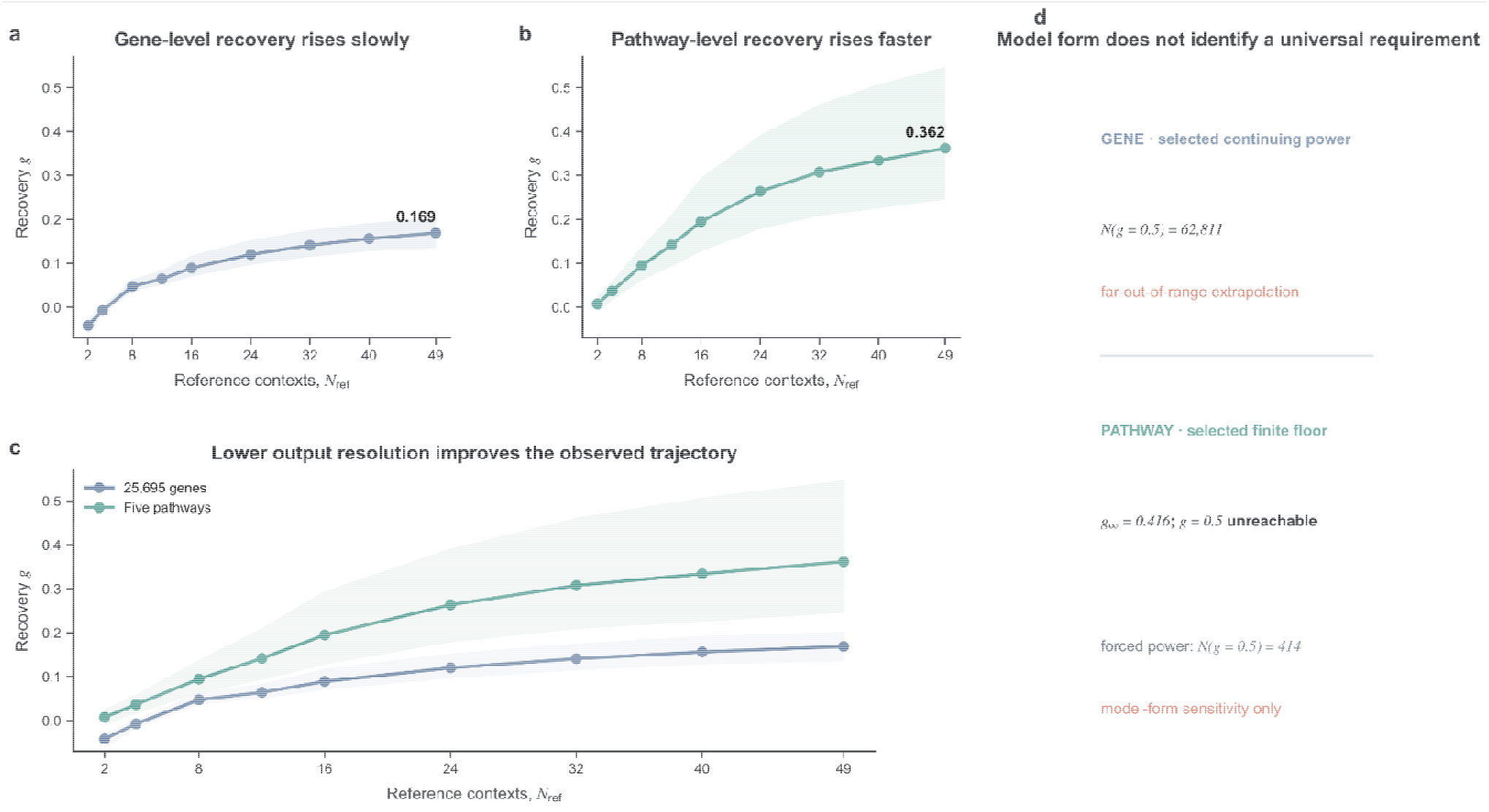
Experimental support and output resolution define distinct scaling boundaries. a,b, Gene- and pathway-level recovery in a held-context scaling analysis, where all target-context responses were withheld and response-observed reference contexts were available. Recovery over the frozen support grid reached 0.168976 (95% percentile interval, 0.134613–0.202541) and 0.361825 (0.245314–0.547051), respectively, at = 49. c, Overlay of the observed gene and pathway trajectories. At = 49, the paired pathway-minus-gene recovery difference was 0.192849 (95% percentile interval, 0.083457–0.388801). Bands in a–c show intervals from the prespecified context-first/intervention-within-context hierarchical bootstrap (10,000 draws), which resampled target contexts and then interventions within each sampled context. d, The selected continuing-power gene fit projects approximately 62,811 response-observed reference contexts, each supplying all 93 interventions, to g=0.5. The selected pathway fit has a finite asymptote of 0.416142; a matched continuing-power sensitivity fit projects g=0.5 at 413.677 such contexts. The extrapolations describe this assay, estimator and representation.

**Extended Data Fig. 8.**
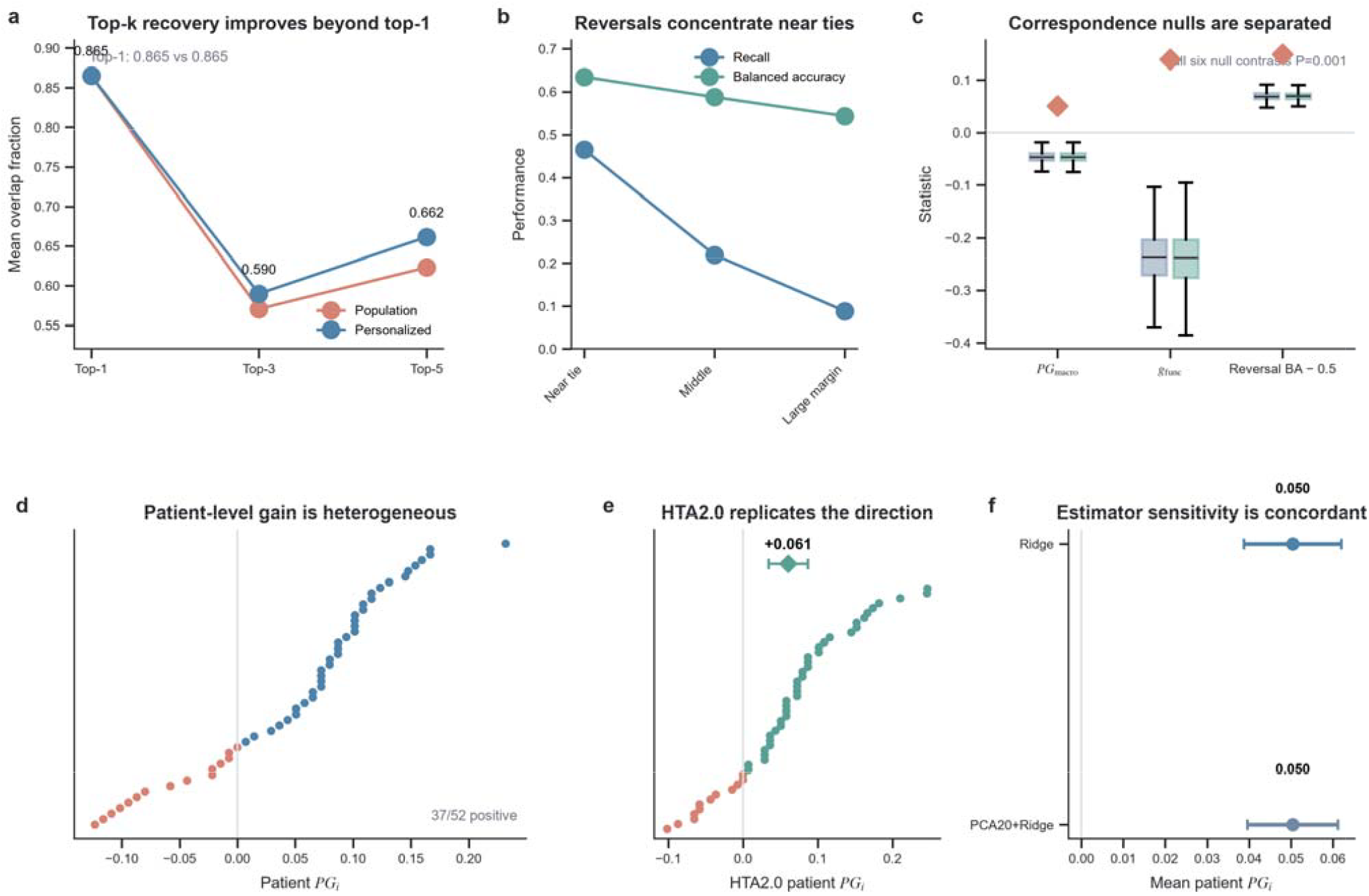
Patient-specific ex vivo drug prioritization is partial, reproducible and bounded. a, Top-1, top-3 and top-5 prioritization under the patient-disjoint RNA model and population reference, respectively (0.865 versus 0.865, 0.590 versus 0.571 and 0.662 versus 0.623). The top-1 near-tie distinguishes stable best-drug selection from broader ordering gains. b, Reversal recall and balanced accuracy across population-margin strata; reversals concentrate among near-tied drug pairs. c, Two within-fold training-patient correspondence nulls shuffled baseline RNA profiles or complete residual-response profiles while preserving drug labels and within-patient response-profile coherence. All six metric-by-null add-one tail probabilities were P=1/1,001; the three-endpoint primary sign-flip family used max-T correction. d, Patient-level personalization gain is heterogeneous, with 37 of 52 patients above zero. e, The secondary HTA2.0 cohort reproduces the positive personalization direction (mean gain, 0.060534; simultaneous 95% interval, 0.034257–0.086810). f, PCA20 plus Ridge gives a concordant positive sensitivity estimate. All evaluations use patient-disjoint folds and ex vivo drug-sensitivity endpoints.

**Extended Data Fig. 9.**
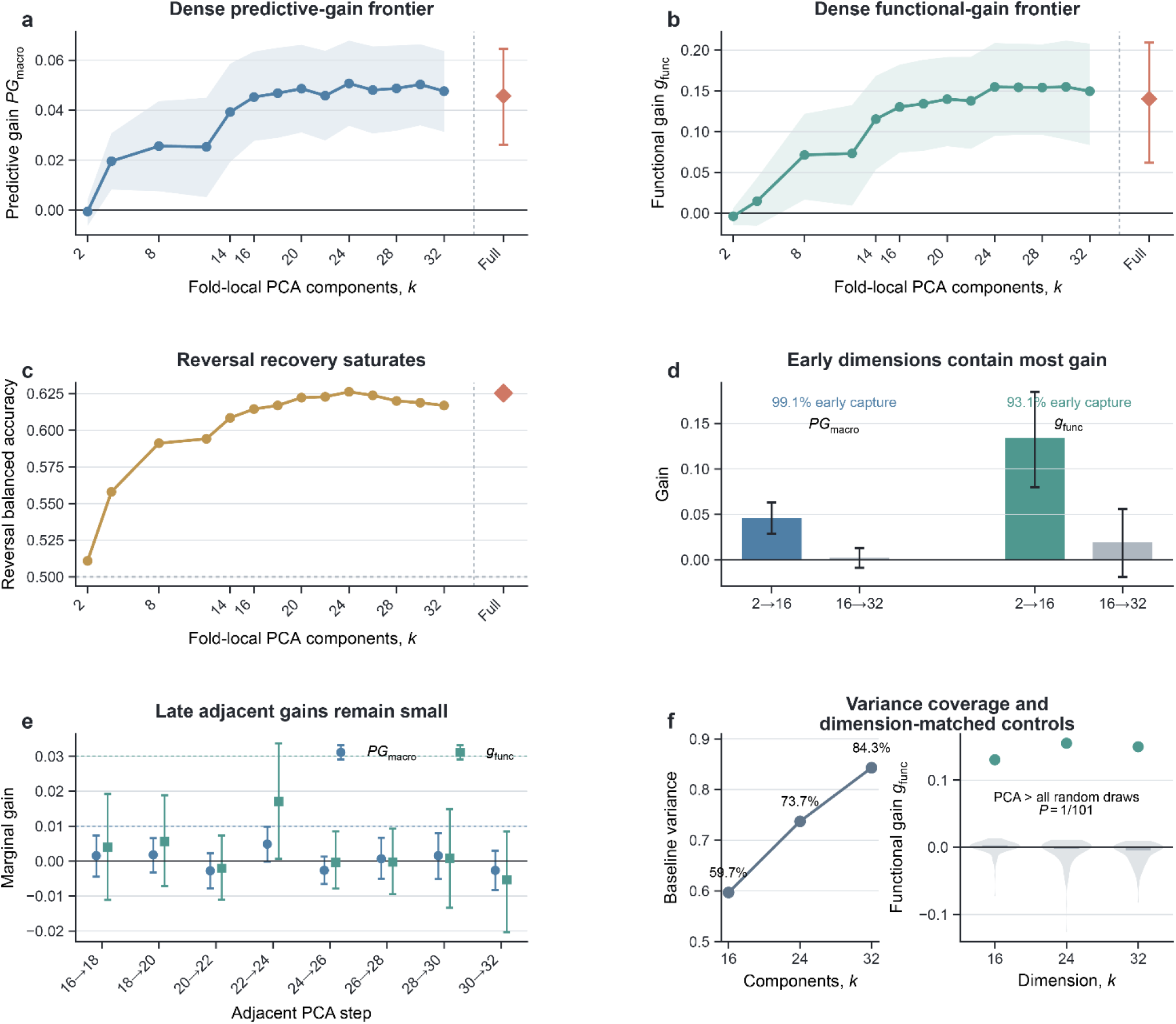
Predictive information saturates across a compact baseline representation range. a,b, Mean personalization gain and functional recovery across the dense fold-local PCA grid (2–32 components) in 52 strictly held patients; shaded bands show patient-bootstrap uncertainty and orange diamonds show the reoptimized full-RNA estimates from 19,421 genes. c, Preference-reversal balanced accuracy across the same representations. d, Gains accumulated from 2 to 16 components (0.045799 for personalization gain and 0.134091 for functional recovery) compared with the much smaller gains from 16 to 32 components (0.002367 and 0.019365). PCA16 captured 99.1% and 93.1% of the corresponding PCA2-to-full-RNA gains. e, Paired adjacent increments through the 16–32-dimensional region; no increment reached the prespecified practical scales of 0.010 for personalization gain or 0.030 for functional recovery. f, Left, training-expression variance explained at PCA16, PCA24 and PCA32 (59.7%, 73.7% and 84.3%). Right, PCA16, PCA24 and PCA32 exceeded all 100 dimension-matched random projections for functional recovery (empirical P = 1/101); the personalization-gain control was concordant. PCA transformations were fitted within training patients only. The plateau summarizes held-patient performance using components derived from full RNA.

