## Supplemental Information for "The recoverable resolution of cellular perturbation-response prediction"

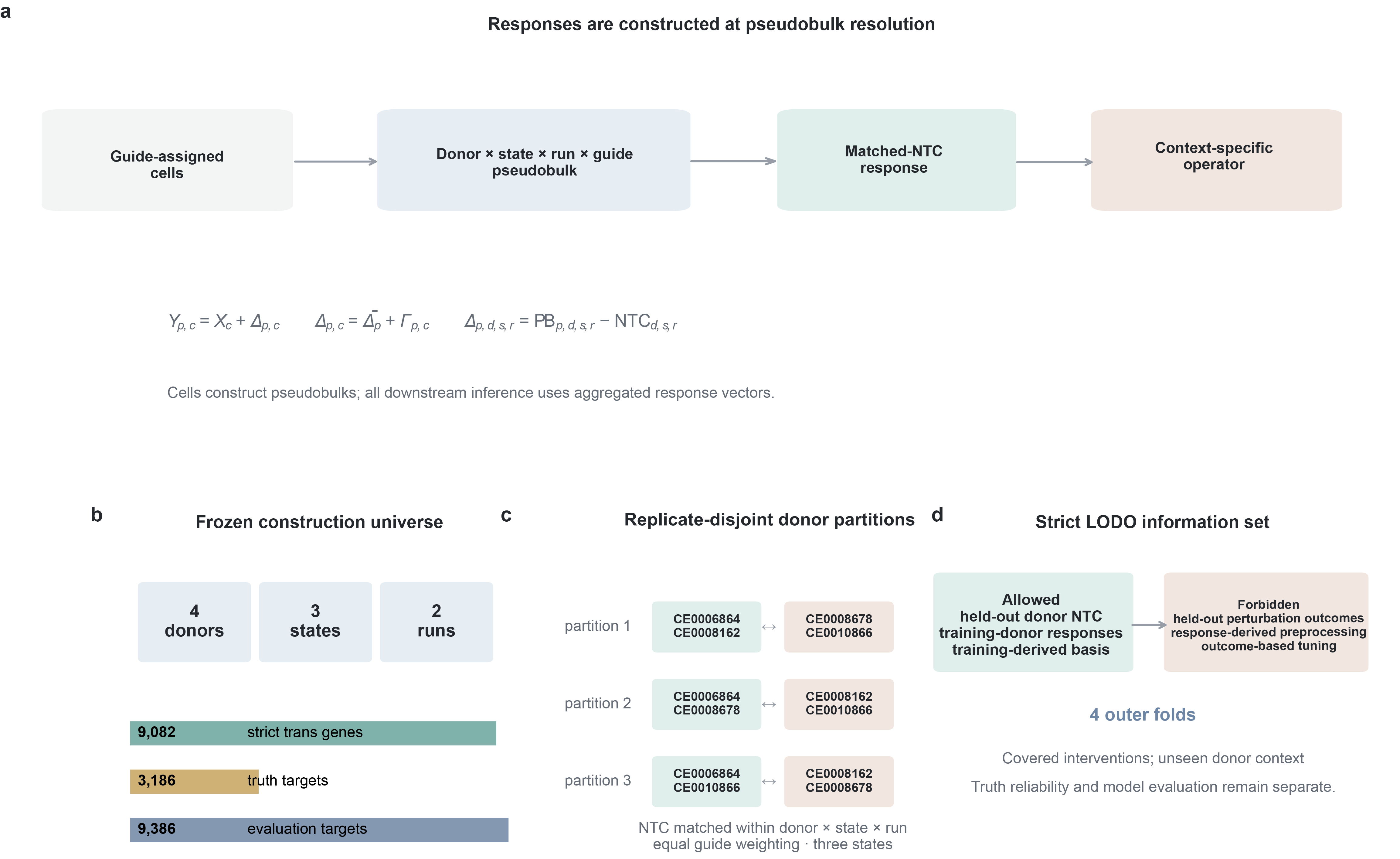

Supplementary Fig. 1 | T-cell pseudobulk construction, guide-disjoint truth and strict held-donor information sets. a, The analysis used author-provided donor × state × run × guide pseudobulks; no per-cell prediction was performed. Perturbation responses were differences from NTC pseudobulks matched within donor, state and run. b, Prespecified construction universe: four donors, three stimulation states, two sequencing runs, 9,082 strict-trans response genes, 3,186 guide-disjoint complete truth targets and 9,386 model-evaluation targets. c, Three balanced partitions of the four donors into non-overlapping pairs for replicate-disjoint truth comparisons. Separately, targeting guides were split into non-overlapping halves; both halves used the same eligible NTC pool, with equal guide weights and NTC matching within donor × state × run. Truth reliability is isolated from model evaluation. d, Strict leave-one-donor-out information set. Each outer fold uses three training donors and permits the untreated NTC state of the held-out donor, while all held-out-donor perturbation responses are forbidden. This is the covered-intervention, unseen-context problem.

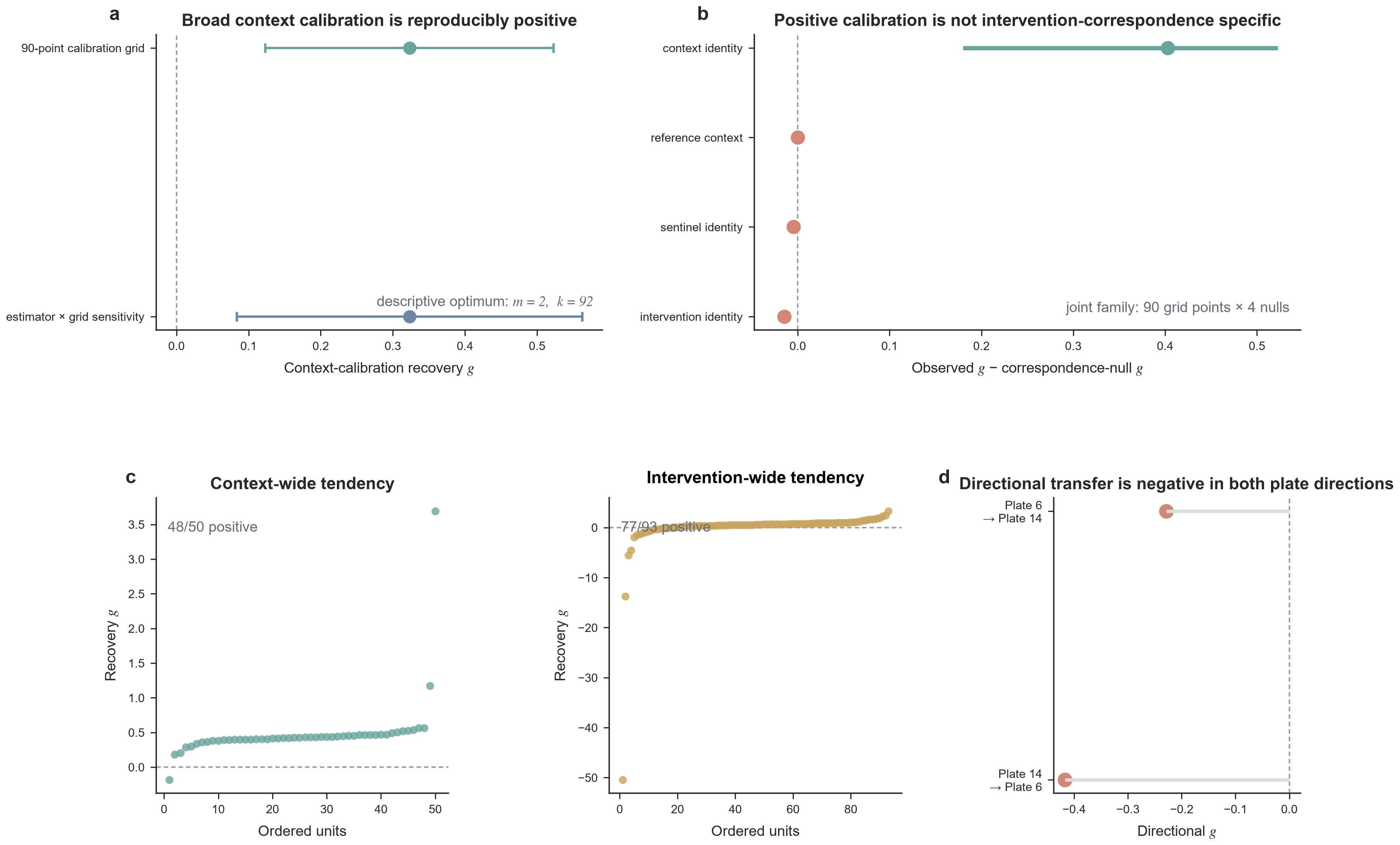

Supplementary Fig. 2 | Broad context calibration is not correspondence-specific recovery. a, The descriptive optimum at two reference contexts and 92 target-context sentinels gives g = 0.323089. Its simultaneous interval is 0.123099–0.523079 over the prespecified 90-point calibration grid and 0.083622–0.562556 under the stricter estimator × grid sensitivity. b, Post-selection observed-minus-null contrasts evaluated jointly over all 90 grid points × four correspondence-null families. The context-identity contrast is 0.402954 (pointwise interval, 0.182295–0.520413; family-wise P = 0.007699); intervention-identity (−0.014703), sentinel-identity (−0.004730) and reference-context (0) contrasts do not pass. Positive pooled association therefore does not establish intervention correspondence. c, Descriptive distributions: 48/50 context estimates and 77/93 intervention estimates are positive, but both contain severe outliers and do not replace the prespecified correspondence test. d, Cross-plate squared-error recovery is negative in both directions (Plate 6 → Plate 14, −0.228463; Plate 14 → Plate 6, −0.417226).

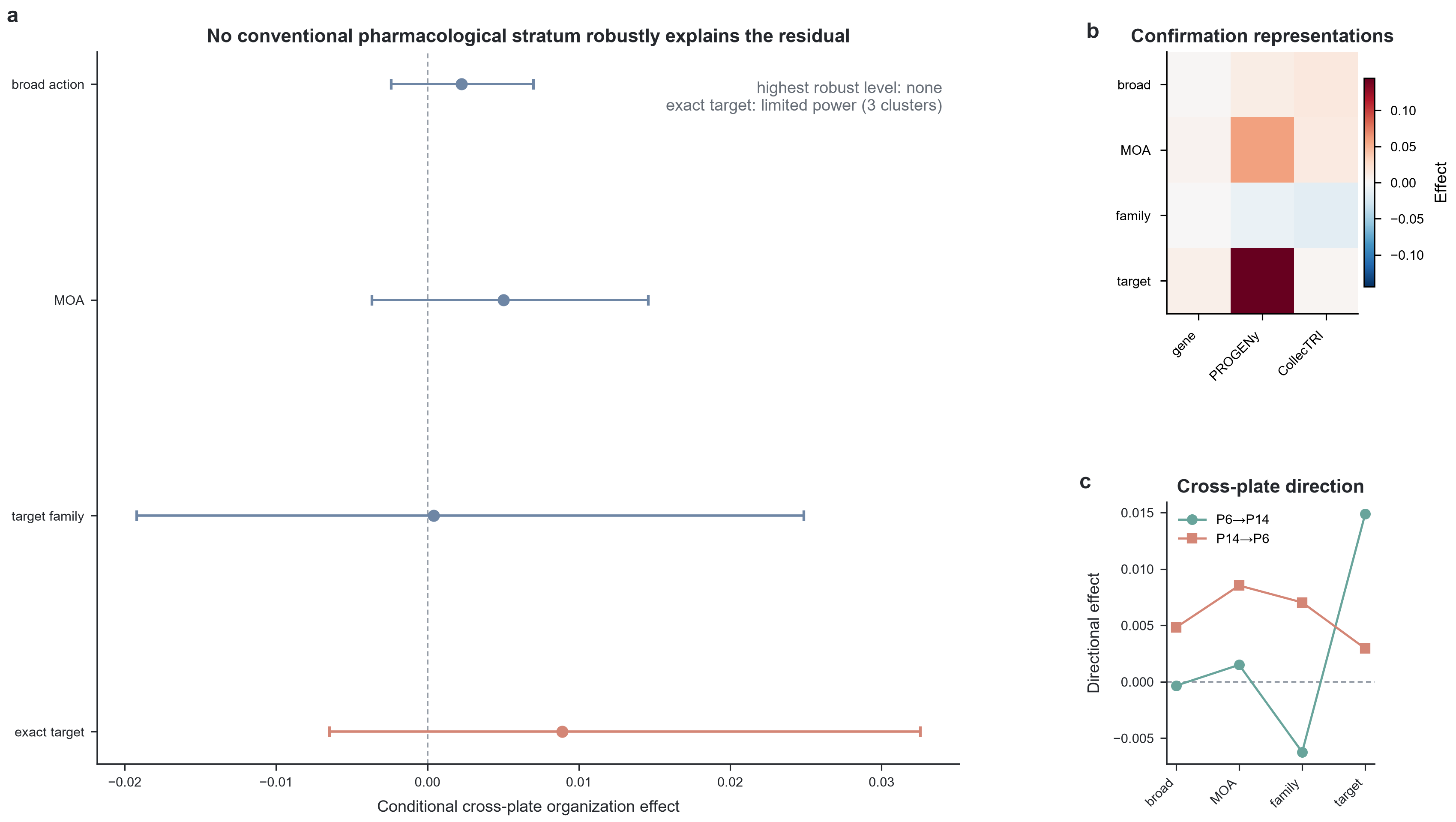

Supplementary Fig. 3 | Conventional pharmacological strata do not robustly explain the unresolved operator. a, Pooled gene-space conditional effects and cluster intervals: broad action, 0.002237 (−0.002383 to 0.007009; two clusters; not estimable); mechanism of action, 0.005016 (−0.003659 to 0.014596; eight clusters; not robust); target family, 0.000399 (−0.019223 to 0.024878; ten clusters; not robust); and exact target, 0.008912 (−0.006483 to 0.032565; three clusters; limited power). No hierarchy level meets the prespecified robustness criterion. b, Pooled effects across gene-space, PROGENy and CollecTRI representations for the same pharmacological hierarchy. c, Cross-plate directional effects for Plate 6 → Plate 14 and Plate 14 → Plate 6. The result bounds the tested conventional pharmacological hierarchy; exact-target evidence is power-limited and is not evidence of absence.

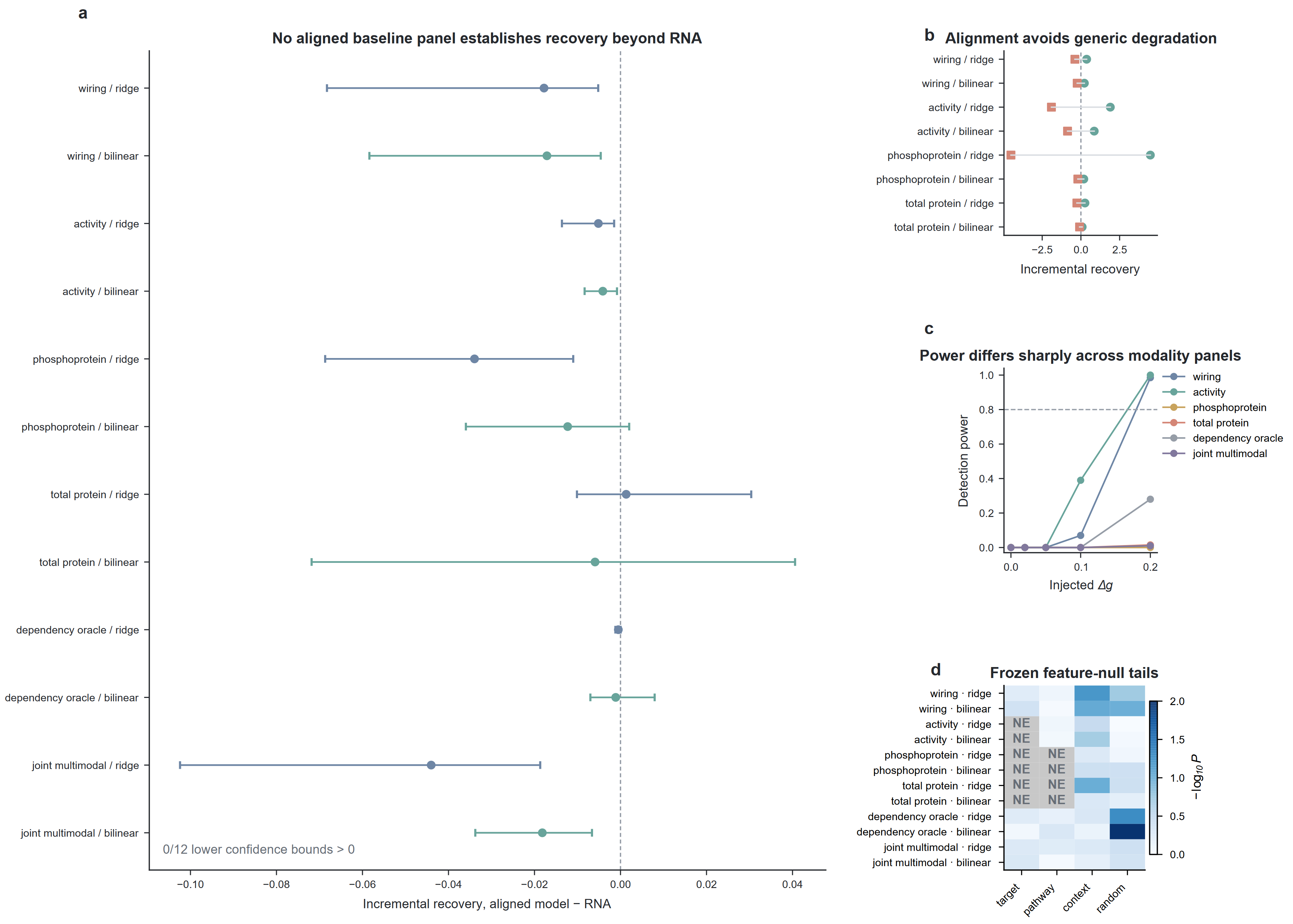

Supplementary Fig. 4 | Mechanism alignment avoids generic-feature degradation but does not establish recovery beyond RNA. a, Twelve prespecified aligned-versus-RNA contrasts across wiring, pathway activity, exact-target phosphoprotein, exact-target total protein, dependency-oracle and joint multimodal panels, each estimated with additive multi-output Ridge and bilinear Ridge. Zero of 12 lower confidence bounds exceeds zero. b, Recovery contrasts for the eight executable ordinary-modality comparisons: teal circles, aligned minus generic-feature models; coral squares, generic-feature minus matched RNA-only models. Alignment often mitigated the degradation produced by generic features. c, Prespecified fold-local power curves. Detection power first reached 80% at an injected oracle-aligned recovery increment of Δg=0.20 for wiring and activity; it did not reach 80% by Δg=0.20 for exact-target phosphoprotein, exact-target total protein, dependency-oracle or joint multimodal panels. These thresholds quantify detection power for injected oracle-aligned signal. d, One-refit feature-null bootstrap tails for four correspondence-null families; non-executable cells are marked NE. One executable contrast interval exceeds zero, but its observed aligned-versus-RNA gain is negative and therefore cannot establish recovery. Panel universes and complete-case information sets are reported in Supplementary Table 6.

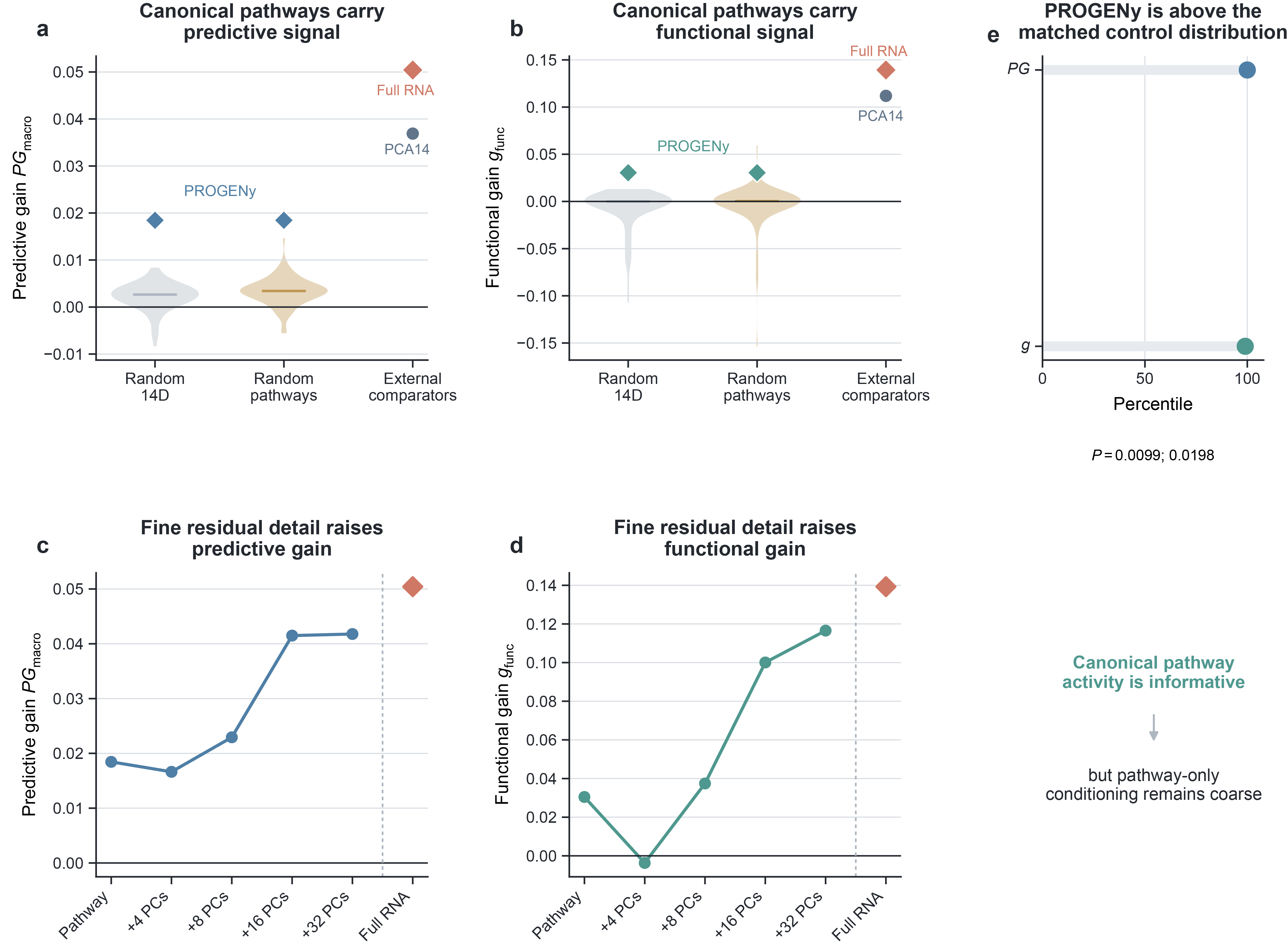

Supplementary Fig. 5 | Canonical pathway state is informative but does not exhaust patient-specific predictive information. a,b, Personalization gain and functional recovery for the rank-14 PROGENy baseline representation relative to 100 14-dimensional random projections, 100 matched random pathway vocabularies, PCA14 and full RNA in the 52-patient, 24-drug leave-one-patient-out analysis. PROGENy 14 achieved ${PG}_{macro}$= 0.018453 and $g_{func}$= 0.030458, compared with 0.036888 and 0.111881 for PCA14 and 0.050423 and 0.139289 for full RNA. c,d, Adding fold-local residual components orthogonal to the PROGENy span increased predictive and functional gain. Relative to pathway-only conditioning, 16 residual components increased personalization gain by 0.023043 (95% interval, 0.007425–0.038661) and functional recovery by 0.069608 (0.010635–0.128580); 32 residual components increased the same endpoints by 0.023329 (0.009725–0.036932) and 0.086095 (0.035367–0.136823). e, PROGENy lay at the upper edge of both matched random-pathway control distributions; empirical P values are shown. All pathway and residual representations were fitted or applied within training patients only. Canonical pathway activity therefore carries real patient-specific signal, but a 14-pathway conditioning representation remains too coarse to preserve the full functional personalization available in baseline RNA.
